# Topology-Driven Negative Sampling Enhances Generalizability in Protein-Protein Interaction Prediction

**DOI:** 10.1101/2024.04.27.591478

**Authors:** Ayan Chatterjee, Babak Ravandi, Parham Haddadi, Naomi H. Philip, Mario Abdelmessih, William R. Mowrey, Piero Ricchiuto, Yupu Liang, Wei Ding, Juan C. Mobarec, Tina Eliassi-Rad

## Abstract

Unraveling the human interactome to uncover disease-specific patterns and discover drug targets hinges on accurate protein-protein interaction (PPI) predictions. However, challenges persist in machine learning (ML) models due to a scarcity of quality hard negative samples, shortcut learning, and limited generalizability to novel proteins. Here, we introduce a novel approach for strategic sampling of protein-protein non-interactions (PPNIs) by leveraging higher-order network characteristics that capture the inherent complementarity-driven mechanisms of PPIs. Next, we introduce UPNA-PPI (Unsupervised Pre-training of Node Attributes tuned for PPI), a high throughput sequence-to-function ML pipeline, integrating unsupervised pretraining in protein representation learning with topological PPNI samples, capable of efficiently screening billions of interactions. UPNA-PPI improves PPI prediction generalizability and interpretability, particularly in identifying potential binding sites locations on amino acid sequences, strengthening the prioritization of screening assays and facilitating the transferability of ML predictions across protein families and homodimers. UPNA-PPI establishes the foundation for a fundamental negative sampling methodology in graph machine learning by integrating insights from network topology.

## INTRODUCTION

Proteins play a central role in essential biological processes, including catalyzing reactions, transporting molecules, responding to pathogens in the immune system, and facilitating cell-to-cell signal transduction [1, 2]. Moreover, crucial cellular processes vital for our health, such as DNA replication, transcription, translation, and transmembrane signal transduction, depend on specific functional proteins [3]. These fundamental biological activities are regulated through protein complexes, typically governed by protein-protein interactions (PPIs) [4, 5]. In humans, deviations from typical patterns of PPIs and protein complexes can either cause or indicate a disease state [6]. Numerous computational methods have been developed to uncover the etiology of diseases form PPIs [7]. However, the incompleteness of the human interactome [8, 9] hinders the understanding of pathogenic and physiologic mechanisms that trigger the onset and progression of diseases [10], and hence in the development of novel therapeutic strategies [11].

Experimental PPI databases such as BioPlex [12], STRING [13], APID [14], BioGRID [15], CoFrac [16], CORUM [17], HuRI [18], HINT [19], and HIPPIE [20], to name a few, capture human PPIs observed via Affinity Purification - Mass Spectrometry (AP-MS) [21] and Yeast-to-hybrid (Y2H) assays [22]. However, none of the PPI databases report the failed experiments, i.e., the non-interactions, creating a scarcity of high-quality protein-protein non-interactions a.k.a. hard negatives [23] in training ML models (see Figure 1A). Negatome [24] stands out as the sole database that has endeavored to tackle this concern; however, it encompasses merely 2,424 interactions linked to human proteins [25] and is limited by specific environmental constraints. As a result, it falls short of capturing the essential interaction mechanisms required for training machine learning models to prioritize large-scale screening. Zhao et al. [26] provided triple-layer validated [27] negatives, yet only 58 pairs are provided for the human proteins reported in UniProt [25]. Furthermore, these databases suffer from high rates of false negatives and false positives [28–30], which combined with the limitations of the traditional ML-based negative sampling methods significantly bias the PPI predictions [31]. For example, some authors propose generating high-quality non-interactions by considering pairs of proteins with distinct cellular localization [32], presumably hindering their participation in biologically relevant interactions [33, 34]. This method samples protein pairs as negative samples from the compartment pairs that do not share any protein (see Figure S1). Alternatively, other authors adopt a simpler approach, randomly selecting protein-protein non-interaction (PPNI) pairs from the entire set of protein pairs that are not known to interact [35–38]. However, both negative sampling scenarios produce a skewed distribution of negative examples resulting in overly optimistic estimates of classifier accuracy [33, 39, 40]. Furthermore, Bardes et al. [41] have shown that the necessity of negative sampling while training semi-supervised models for predicting similarity between two entities can be overcome by regularization. Nevertheless, as the similarity of proteins does not necessarily indicate interaction, the need for negative sampling persists in PPI prediction. The complementary nature of the PPI network [42] demands hard non-interaction samples for unbiased prediction and improved generalizability. In a recent paper, Li et al. [43] have proposed a heuristic-based hard-negative generation method for link prediction. However, this method lacks biological and chemical rationality in creating protein-protein non-interactions and impedes the interpretability of the predictions. Finally, the complex interdependence of the in-vitro screening experiments on the environmental factors makes it difficult to obtain hard negatives solely driven by the molecular properties of the proteins [44]. In another recent paper, diffusion models have been used for deriving quality negative samples for link predictions [45]. Understanding the evolutionary reasoning for protein interactions and non-interactions is a complex task. However, recent research has unveiled the complementarity-driven mechanisms [42] driving protein-protein interactions, which include an electrostatic charge or Coulombian complementarity [46], hydrophobic mismatch [47], conformational complementarity [48], hydrogen bond complementarity [49], etc. (see Figure 1C). Therefore, delineating interactions and non-interactions based on the complementary nature of the PPI network not only facilitates the generation of hard negative samples for training PPI prediction models but also enhances the mechanistic interpretability of these predictions.

**Figure 1:**
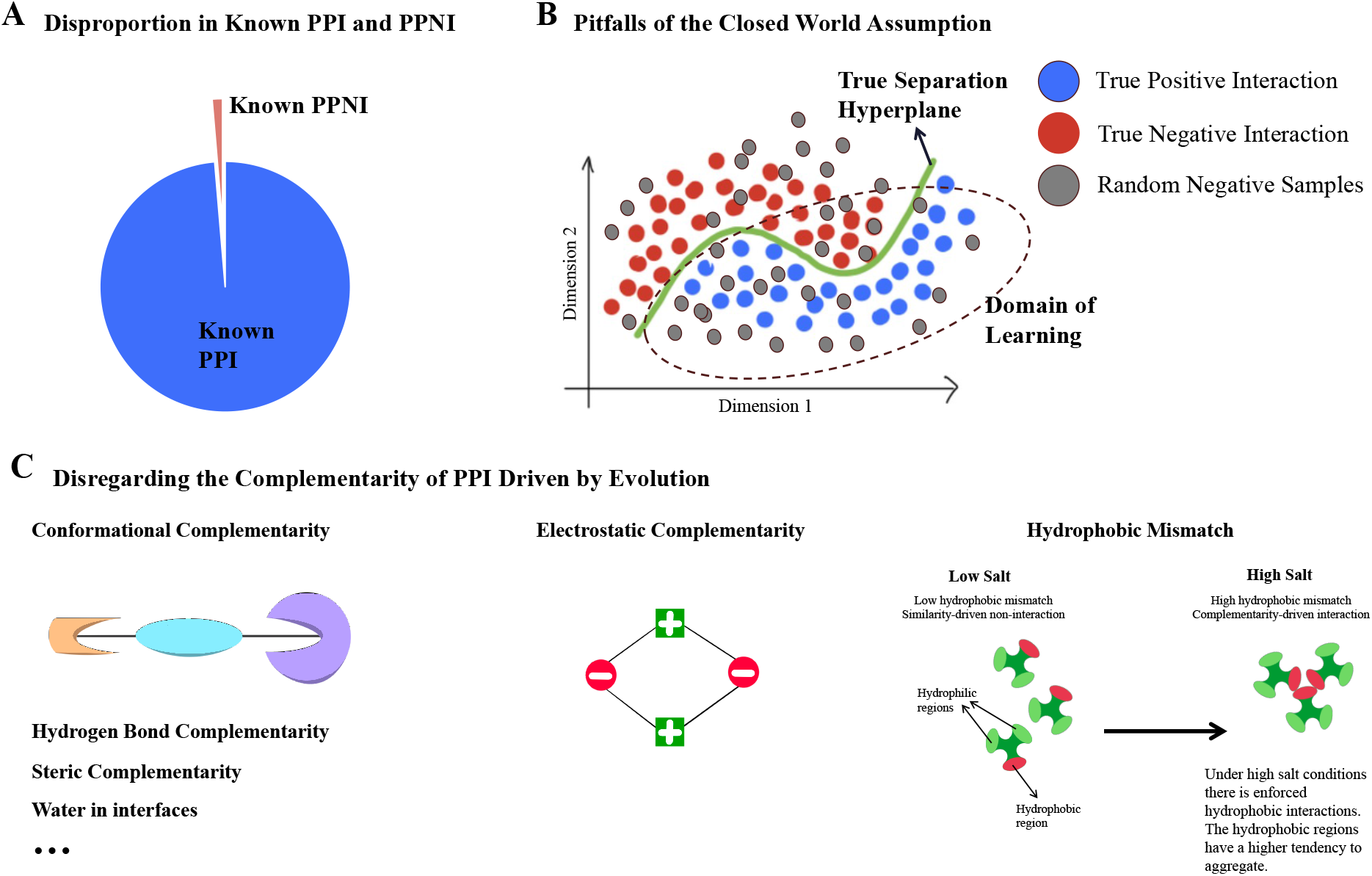
Status quo in machine learning-based PPI prediction. **(a)** We observe a substantially higher number of PPI samples in the existing databases compared to PPNI samples. The lack of true negative samples is a major obstacle in training reliable machine learning models for PPI prediction. **(b)** Traditional machine learning training methodology utilizes random sampling from the complement graph (G^*c*^) of the training PPI network (G) to obtain negative examples. However, this approach violates the closed-world assumption, i.e., not observing a PPI does not imply non-interaction between those proteins. Hence, the traditional approach fails to create high-quality PPNI and shifts the domain of learning in the loss function manifold, hampering the machine learning models from learning the true separation hyperplane between PPI and true PPNI. **(c)** In similarity-driven networks, entities that are alike are connected by links. For example, in social networks, people with similar interests are connected. On the other hand, in complementarity-driven networks, entities with opposing properties are linked. We list six complementary properties of proteins that are major contributors to protein-protein interactions [49]. Leveraging the complementary nature of PPI networks has recently gained much attention and bifurcates the machine learning approaches for PPI prediction from those widely used in social network analysis.

Historically, the scarcity of hard negatives and the pronounced imbalance between edges and non-edges during the training of machine learning models for link prediction tasks have led to the practice of obtaining negative samples through random sampling from the complementary graph [50]. Nevertheless, it is noteworthy that random negative samples can introduce substantial bias into the separation hyperplane learned by the classifier, as illustrated in Figure 1B. Selecting an appropriate set of hard negatives is essential for acquiring the relevant hyperplane and enhancing generalizability in machine learning [23].

On a separate note, observation bias [51, 52] and shortcut learning [53, 54] are prevailing in the ML-based PPI prediction models due to selection and laboratory biases in the PPI databases [55]. The excellent transductive (see Section S1, Figures S2 and S3) cross-validation performances of the majority of the state-of-the-art (SOTA) PPI prediction models [56, 57], falsely exaggerated by topological shortcuts [52,54,58], are often misleading and not biologically interpretable. Multiple recent papers have focused on the poor performance of state-of-the-art link prediction models on low-degree nodes [43,51,59]. GraphPatcher [59] proposes a novel test-time augmentation method to improve link prediction performance for the low-degree nodes.

However, the approach is infeasible for unseen nodes, making the prediction task for never-before-seen nodes of greater difficulty [51], which maps to the scenario of predicting interactions between novel proteins. Therefore, inductive tests have garnered significant attention recently for their capability to reveal the authentic predictive efficacy of machine learning models (see Figure S2). This allows for the assessment of a model’s performance on novel entities, ultimately contributing to enhanced generalizability and interpretability [54, 60]. Recent studies [60, 61] emphasize the importance of going beyond the PPI network topology in machine learning and advocate for the incorporation of inductive tests in constructing models with biological utility and interpretability. Yet, intricate calibration of these models via fine-tuning [62] and embedding regularization limit their generalizability within particular databases and families of proteins [60].

Finally, using the appropriate performance metric is essential for evaluating the true performance of an ML model [63, 64]. Global performance metrics like precision, recall, the area under the receiver-operating characteristics (AUROC), the area under the precision-recall curve (AUPRC), and accuracy fail to capture the local performance in the PPI network [65]. Hence, performance metrics like Hits@TopK [66] and mean reciprocal rank (MRR) [67], which incorporates local evaluation [68], are necessary for assessing an ML model for novel, unseen proteins in inductive tests. Furthermore, the evaluation of negative predictions is often overlooked. However, the ability of an ML model to distinguish the positives from the negatives largely determines its predictive power [69]. Yet, a systematic approach is absent in evaluating PPI prediction models.

The **contributions** of this work are as follows:

1. We introduce UPNA-PPI (Unsupervised Pre-training of Node Attributes tuned for PPI), an ML pipeline that bridges the existing gaps in the PPI prediction methodology by
  a. including a method for obtaining high-quality protein-protein non-interaction (PPNI) pairs, which utilizes the PPI network topology;
  b. connecting PPNI pairs to complementary generation mechanisms through hyperbolic embedding space;
  c. improving inductive link prediction performance in PPI using topological negatives; and
  d. enabling interpretability and transferability of PPI prediction across protein families and homodimers.
2. We propose local performance metrics to evaluate both interaction and non-interaction predictions, quantifying the distinguishing power of any PPI prediction model.
3. We predict and validate interactions between understudied and difficult-to-purify G protein-coupled receptors (GPCRs).

## RESULTS

### Topological PPNI

Negative sampling is an inextricable part of binary classification task in machine learning [70]. The lack of high-quality hard negatives in biological data is a major hindrance in developing practical ML models. The majority of the biological databases have a sufficient number of positive examples, but a considerably low number of negative samples due to experimental and observational biases [52], which is a major cause of both overall class imbalance and node-wise class imbalance (annotation imbalance) [54]. Random negative sampling is the most frequently used approach to mitigate this problem. Yet, random negative samples are consequential in closed systems [71] and are prone to degree bias [54, 59]. In this section, we propose a novel method of creating hard PPNIs by leveraging the PPI network topology [72], which is underpinned by the complementary nature of the interactome [42]. We propose topological PPNI (see Figure 2A) that combines entropy-based network null models [73] and higher-order PPI network properties [72] in prioritizing hard PPNI samples. Topological PPNI methodology consists of two parts: **(a)** a traditional unipartite configuration model, and **(b)** contrastive-L3 (CL3) filtering.

**Figure 2:**
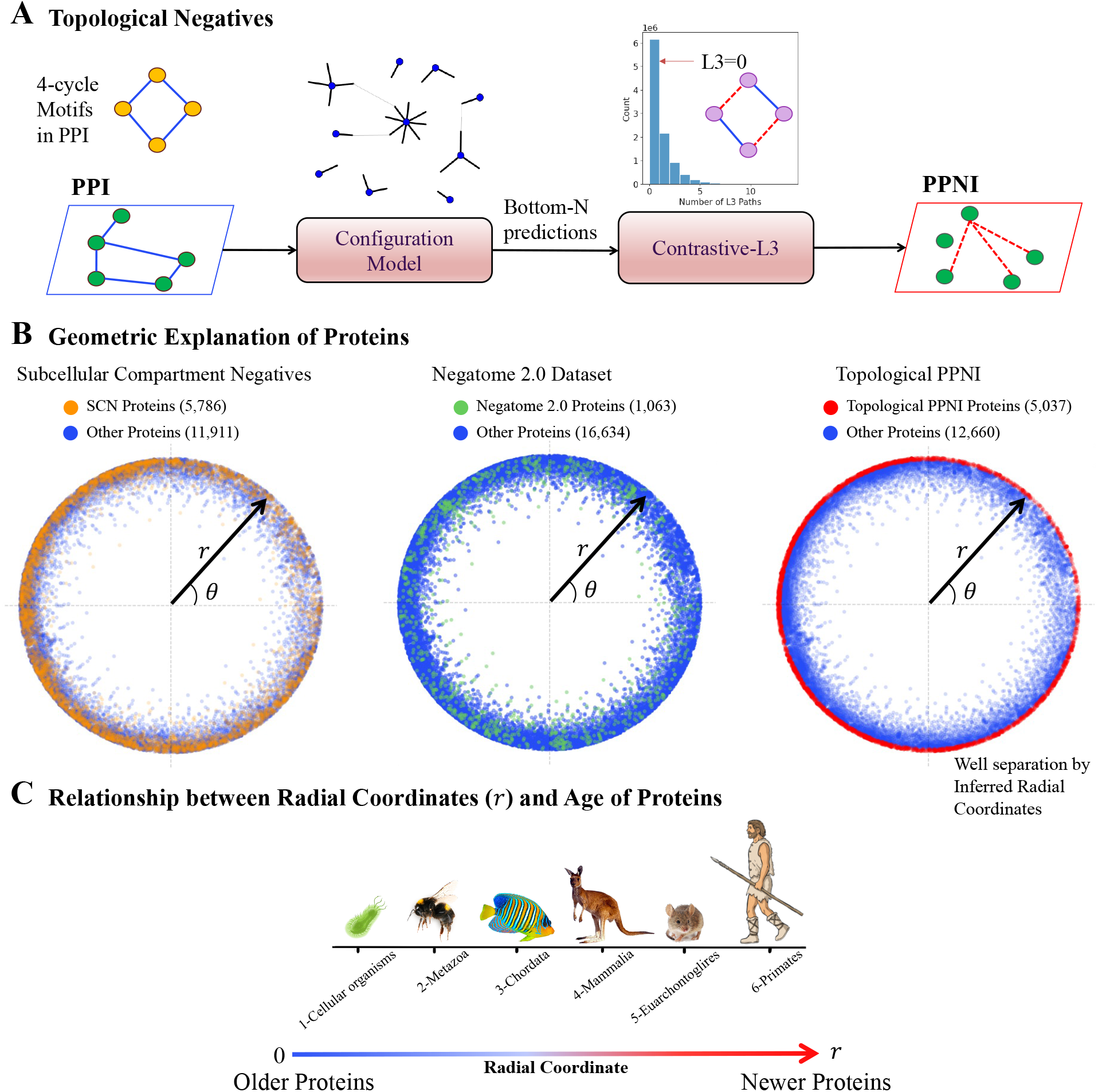
Leveraging PPI topology for PPNI sampling. **(A)** We propose a novel method for sampling high-quality PPNI leveraging the topology of PPI driven by its complementary nature. First, we run a traditional configuration model to identify the topologically least probable edges via entropy maximization. The bottom-N edges are then used for computing the 4-cycles or L3 paths induced by the protein pairs. (We use N=10 million.) The configuration model helps in reducing the protein-pair search space from 156M down to 10M. Interacting pairs induce many L3 paths in the PPI network. We utilize the inverse hypothesis, namely *Contrastive L3 or CL3*, to filter the protein pairs from the bottom-N predictions which induce no L3 path in the PPI network. Hence, we obtain the topological PPNI, which is used in the training and testing of UPNA-PPI. **(B)** Hyperbolic embeddings have recently been used for visualizing and link prediction tasks in complementarity-driven networks like PPI networks. We visualize the proteins involved in PPI and different PPNIs in the hyperbolic space. We observe that the proteins involved in subcellular compartmental negatives (SCN) and PPNI from the Negatome database show a large overlap with proteins in the PPI. However, the proteins involved in topological PPNI show a clear separability from the proteins in PPI. Proteins involved in topological PPI are towards the circumference of the hyperbolic disc and are hence constituted by evolutionarily younger proteins. **(C)** Alanis-Lobato et al. [75] established the relationship between the radial coordinates ***r*** of human proteins and their age by assigning proteins to six different age groups through grouping proteins based on their ancient relatives in other species (subfigure courtesy of Alanis-Lobato et al. [75]). The clear selection of younger proteins by *CL3* hypothesis indicates that *CL3* can only identify negative interactions for younger proteins, which are less central in biological pathways. Hence biology would optimize less competition between the complementarity mechanisms to form a biologically relevant function for younger proteins, leaving patterns in PPI topology that can distinguish younger and older proteins as captured by *CL3*.

We use the traditional unipartite configuration model (see Methods), which takes as input only the degree sequence of the PPI network and achieves a commendable performance in transductive link prediction (AUROC: 0.87 ± 0.0002, AUPRC: 0.87 ± 0.0002, Hits@Top100:^*^ 0.98±0.016). The output of the configuration model dictates the interaction probability between two proteins from a topological standpoint. We select the bottom-N (N=10 million) predictions from this configuration model on all human protein pairs and consider them as less probable for binding, a.k.a, potential hard negative samples. Next, we use a complementarity-driven topological property of the PPI network to derive the hard negatives. Unlike the formation of triadic closure in social networks [74], proteins tend to form quadratic closures or even-length cycles [72]. Two friends in a social network have multiple common friends and hence, many paths of length 2 (L2). On the other hand, two interacting proteins have many paths of length 3 or L3 between them, which is related to the evolutionary aspects and the complementary nature of PPIs [42,72]. We propose the Contrastive-L3 (CL3) hypothesis (see Figure 2A), which states that two non-interacting proteins would not constitute any odd-length (L=3) path in the PPI. We now use CL3 hypothesis to filter high-quality hard negatives from the output of the configuration model. We filter the output of the configuration model mentioned above to fetch the protein pairs that induce no L3 path in the PPI and obtain 3,063,605 hard negatives (see Figure 2A). Note that, ∼18,000 human proteins in different PPI databases can computationally create ∼156 million potential pairs. This is a computationally infeasible space for computing all L3 paths. Therefore, adding the non-interaction prediction from the configuration model significantly reduces the L3 computation space and provides a regime consisting of proteins lacking sufficient annotations, constituting the potential space for sampling hard PPNIs.

### Geometric and Evolutionary Relevance of PPNI

Recent studies have demonstrated the ability of hyperbolic embeddings to capture evolutionary patterns driven by complementarity in PPIs [75]. Figure 2 illustrates various PPNI sampling methods in a two-dimensional hyperbolic space [75, 76]. We use the NetHypGeom hyperbolic embedder proposed by Alanis-Lobato et al. [75]. In Figure 2B, we visualize proteins involved in Subcellular Compartment Negatives [33, 34] (SCN, see Fig S1), Negatome 2.0 [24], and topological PPNIs involving distinct proteins. Alanis-Lobato et al. [75] have associated the radial and angular coordinates of the 2D hyperbolic plane with the evolutionary aspects of human proteins. According to their findings, proteins closer to the origin are evolutionarily older, while those with higher radial coordinates are younger (see Figure 2C). We observe that SCN PPNI involves both old and new proteins due to constraints related to subcellular localization. Negatome PPNI is primarily constrained by well-studied older proteins. In contrast, topological PPNI is constrained to younger proteins. From an evolutionary standpoint biological pathways have evolved around older proteins, resulting in their interactions with the majority of other proteins [75, 77]. Conversely, younger proteins exhibit fewer interactions among themselves. Moreover, protein families are linked to angular coordinates within the hyperbolic space, as indicated by research on latent geometry in PPNI networks [75]. The topological PPNI encompasses the entire range of angular coordinates, thereby encompassing samples from diverse human protein families, as illustrated in Figure 2B.

In Figure 3A, we present a visualization of PPI and PPNI edges in a 2D hyperbolic space. The computation of the edge (both PPI and PPNI) embedding between two proteins involves averaging the hyperbolic embeddings of the respective end proteins across each dimension.

**Figure 3:**
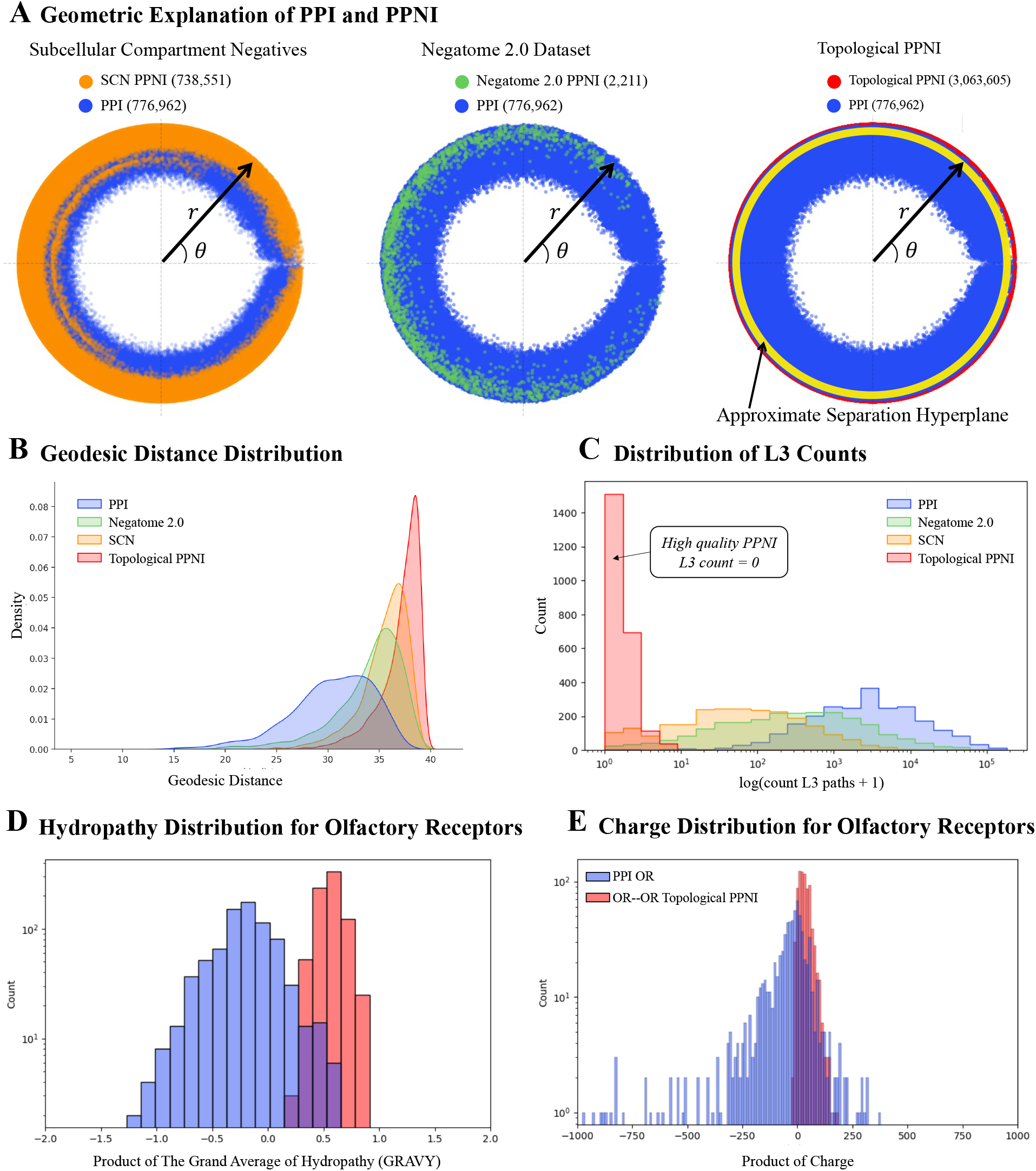
Geometric and Chemical Relevance of topological PPNI. **(A)** We visualize each PPI and PPNI in in the hyperbolic space by averaging ***r*** and θ for each protein pair. Topological PPNI provides the best-separating hyperplane that can be learned by downstream tasks only from the secondary structure of proteins. **(B)** We plot the distributions of geodesic distances between the protein pairs in PPI and different PPNIs (number of samples for each type n=2,311). Topological PPNI identifies pairs of proteins that are significantly far from each other in the hyperbolic geometry of the human PPI network. Hence, compared to SCN and Negatome, topological PPNI offers negative pairs with unique patterns allowing ML to decode and learn such patterns. **(C)** We plot the L3 count distributions for PPI and various PPNIs. While the protein pairs in PPNI induce a significantly lower number of L3 paths in the PPI network, we select the pairs inducing no L3 path in the topological PPNI for training and testing of UPNA-PPI. **(D)** The Grand Average of Hydropathy (GRAVY) is a measure of the hydrophobicity or hydrophilicity of a peptide or protein [82]. The more negative the score, the more hydrophilic the amino acid is, and the more positive the score, the more hydrophobic. Humans have approximately 400 olfactory receptors (ORs) and we observed 7,076 negative inter-family links among ORs in topological PPNI. Also, we have 773 positive links between ORs and other proteins in our PPI, none of the 773 PPI OR links are inter-family. The distribution of the product of GRAVY scores between PPI ORs and OR–OR PPNI (randomly sampled 773 topological PPNI) shows that the *CL3* resulted in identifying pairs of ORs that show higher hydrophobic mismatch, and hence decrease the chance of interacting with each other. **(E)** Similarly, we calculated the product of charge for the 773 PPI OR links and OR– OR topological PPNI (charge at pH = 7). The distribution of the product of charge shows that *CL3* identified inter-family OR proteins with the same sign of charge hence repelling each other and decreasing the chance of interaction.

The substantial overlap observed between SCN PPNI and the experimental PPI implies that SCN fails to effectively capture the complementarity mechanism influencing protein interactions [42]. Consequently, SCN PPNI proves inadequate for training a classifier capable of distinguishing between protein interactions and non-interactions based on complementarity. A parallel observation is made for Negatome PPNI. Conversely, topological PPNI demonstrates effective separation between PPI and PPNI in the hyperbolic space, successfully capturing complementarity-driven interaction mechanisms through negative samples. The hyperplane learned by UPNA-PPI, visualized in Figure 3A, effectively distinguishes PPI from PPNI in the hyperbolic space. While training a classifier on the hyperbolic embeddings would limit the complementarity-driven link prediction to the transductive scenario and would not be able to make PPI predictions for new proteins, UPNA-PPI demonstrates the ability to learn complementary mechanisms of protein interactions from the amino acid sequences when trained using experimental PPI and topological PPNI, and can generalize the learning to new, unseen proteins in inductive tests, hence improving both generalizability and biological interpretability of PPI prediction.

In Figure 3B, we visualize the distributions of pairwise geodesic distances [42] between protein pairs for PPI and various PPNIs. Notably, the pairwise distance distribution for topological PPNI is distinctly separated and positioned to the right of the distribution for PPIs. This indicates that, in the hyperbolic space, interacting proteins are geodesically closer compared to those in topological PPNI. Moreover, topological PPNI demonstrates superior separability between non-interacting proteins compared to SCN and Negatome.

Figure 3C illustrates the L3 paths generated by pairs in PPI and various PPNI scenarios. The visual representation reveals that both SCN and Negatome PPNI induce a significantly lower number of L3 paths in the PPI network when compared to experimentally validated PPIs. To identify topological PPNI, we focus on protein pairs that do not induce any L3 paths in the PPI network. Significantly, given that both L3 paths and the hyperbolic space capture complementarity-driven mechanisms in PPI, protein pairs in the topological PPNI category, which induce no L3 paths, are positioned farthest away in the hyperbolic space. This distinct positioning underscores their clear separation from PPI pairs that exhibit the highest number of induced L3 paths, providing a distinctive characterization.

Finally, we explore a case study involving human proteins to illustrate how topological negatives effectively capture complementary interaction mechanisms. Canonical Olfactory Receptors (ORs) represent a class of G-protein-coupled receptors (GPCRs) in mammals [78]. Monomeric ORs in mammals are activated by chemical ligands and couple to specific G-proteins. For example, ORs in the olfactory sensory epithelium activate Cyclic adenosine monophosphate (cAMP) and other second messenger signaling, leading to ion channel opening and membrane depolarization. Contrastingly, insect ORs function as heteromeric ion channels and lack homology to G-protein-coupled chemosensory receptors found in vertebrates [79, 80]. Humans possess more than 400 ORs, while rodents have approximately 1000 OR genes [81]. Hence, it is biologically justified to observe a higher number of non-interactions between human OR proteins to other human proteins, which is reflected in our topological PPNI samples. We observe a total of ∼15,000 topological negatives involving the human OR proteins, among which ∼7,000 samples include proteins from inter-OR families. Furthermore, biological justification suggests that topological PPNIs are dependent on the species under consideration. Finally, we validate the association between OR-relevant topological PPNI and complementarity mechanisms. In Figures 3D and 3E, we visualize the distribution of the product of hydropathy [82, 83] and electrostatic charge [83] for human OR protein pairs in both PPI and topological PPNI. Divergent distributions, with the PPNI distributions being more positive, validate that topological PPNI effectively captures hydrophobic mismatch and eletrostatic complementarity mechanisms [42].

### UPNA-PPI

Now we use the topological negatives to design UPNA-PPI, a generalizable, robust, interpretable, and transferable PPI prediction ML pipeline. UPNA-PPI uses unsupervised pretraining of node attributes for improved generalizability to unseen proteins [51].

### UPNA-PPI Architecture

UPNA-PPI uses pair-wise learning [84] combined with protein representations pre-trained in an unsupervised fashion [51] for improving the generalizability of ML-based PPI prediction. Figure 4A visualizes the neural architecture of UPNA-PPI. We use ProtVec [85] for embedding the protein amino acid sequences. ProtVec is trained on ∼ 0.5 million amino acid sequences (1.6 million trigram sequences) available in SwissProt [86].

**Figure 4:**
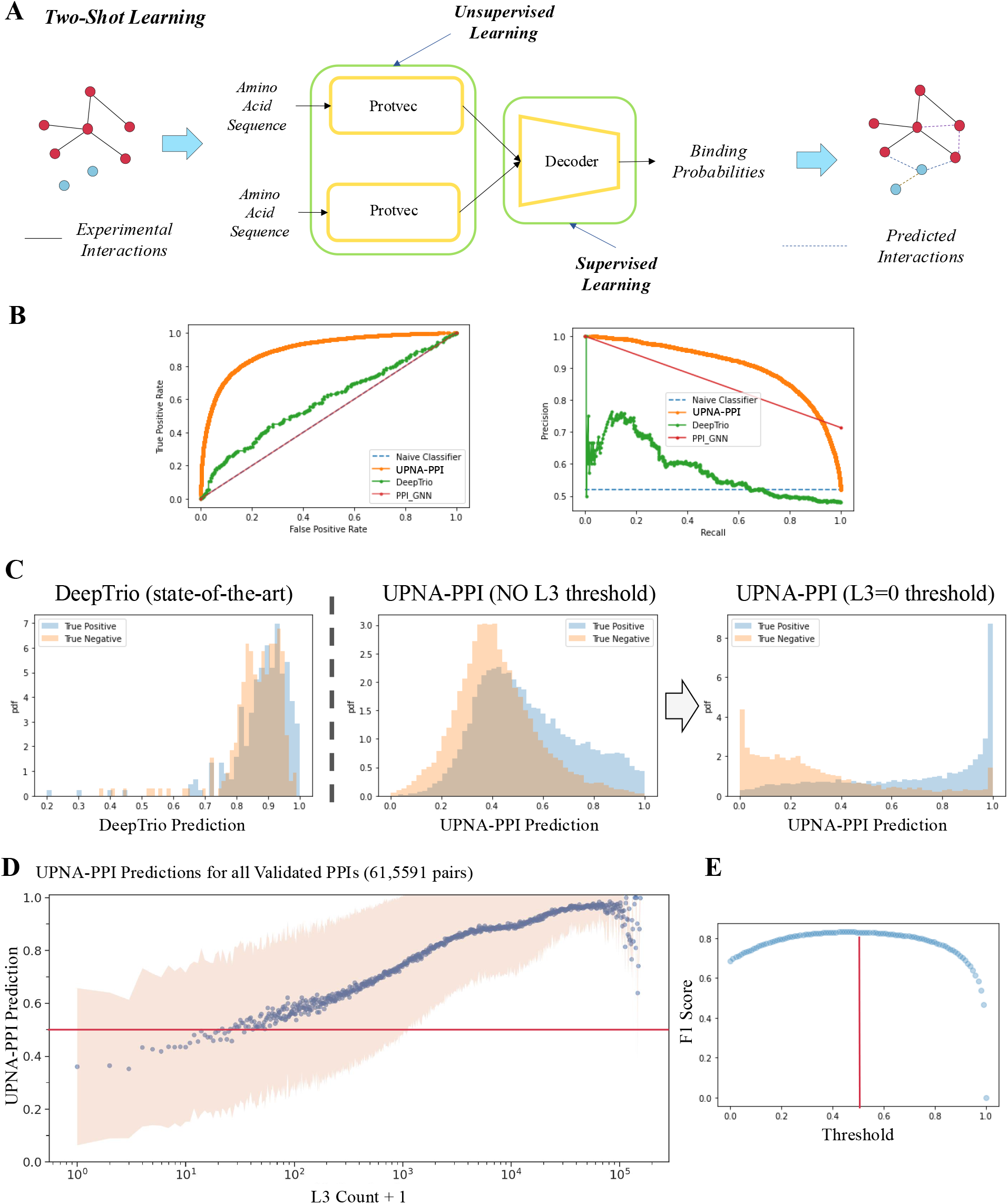
UPNA-PPI architecture, inductive link prediction, and transfer learning. **(A)** UPNA-PPI architecture: UPNA-PPI embeds the protein amino acid sequences into 100-dimensional vectors using ProtVec. For each protein pair, UPNA-PPI concatenates the ProtVec embeddings and feeds it to a decoder (multi-layer perceptron). **(B)** UPNA-PPI achieves better performance both in terms of receiver operating characteristics and precision-recall compared to two state-of-the-art models PPI-GNN and DeepTrio in inductive link prediction. **(C)** DeepTrio predicts overlapping prediction values for the PPI and PPNI test samples. UPNA-PPI shows better separation between the predictions for PPIs and PPNIs, becoming a superior ranking tool for both PPI and PPNI. **(C)** Predictions at the output of DeepTrio are overlapping for the PPI and PPNI data. Hence, DeepTrio is unable to separate the predictions from PPIs vs. PPNIs. UPNA-PPI trained on potential negatives from the configuration model (no *CL3*) shows better separation between the predictions for PPIs vs. PPNIs. Finally, after introducing CL3 thresholding, UPNA-PPI shows a clear separation between the predictions for PPIs and PPNIs. **(D)** We computed the number of L3 simple paths between all PPI pairs to investigate if UPNA-PPI has learned the complementarity mechanisms from protein sequences. Indeed, we observe a strong Spearman’s rank correlation coefficient of 0.48 between L3 counts and UPNA-PPI predictions, indicating that the topological PPNI enforced UPNA-PPI to learn complementary mechanisms that drive protein-protein interactions only from the amino acid sequences. **(E)** We plot the F1-score on the test dataset (first fold) by changing the binary classification threshold. We observe that the test F1-score is maximized for an optimal classification threshold ≈ 0.5. Furthermore, in subplot (D), we observe that UPNA-PPI predicts interaction probabilities greater than the optimal interaction threshold for the majority of the experimentally validated PPIs. Similar observations are made for other folds of UPNA-PPI and the average optimal threshold is 0.476 ± 0.049.

Although machine learning (ML)-based protein-protein interaction (PPI) prediction has been investigated using 3D protein structures [87, 88], recent insights into the dynamic and self-organizing nature of the cell criticize approaches that consider the rigid structures of proteins [89]. The limitations of the current ML PPI prediction models, stemming from the scarcity of experimental 3D human protein structures and the uncertainties associated with AlphaFold [90], impede achieving high accuracy, robustness, and generalizability. The assumed rigidity of 3D structures fundamentally constrains the predictive power of PPI prediction models. Therefore, we opt to utilize amino acid sequences as a basis for learning PPI from protein representations, aiming to overcome these limitations and enhance the predictive capabilities [87–90].

The ProtVec embeddings are fed into two arms of the downstream UPNA-PPI decoder, which formulates a binary classification task. We use a 3-layer multi-layer perceptron (MLP) [91] as the decoder. UPNA-PPI is trained in an inductive setting (see Figure S2) [51]. We use the GoldPPI (see Methods) and Negatome PPNI data in the validation and test datasets for gaining uncompromising confidence in UPNA-PPI predictions through harder tests. Following the inductive link prediction setup from GraIL [92], the human proteins are randomly divided into three groups for train, validation, and testing, with the proteins from the GoldPPI and Negatome PPNI predominantly residing in the validation and test groups. Then, the interactions induced by these proteins are sampled to create the train, validation, and test datasets. We implemented an early-stopping [93] on the inductive validation dataset to avoid overfitting. UPNA-PPI is trained and tested in a 5-fold cross-validation setting.

### Inductive Link Prediction Performance

Two state-of-the-art PPI prediction models, DeepTrio [94] and PPI-GNN [56] perform exceptionally well in transductive link prediction (see Section S2). Yet, their performance significantly diminishes in inductive tests (see Table 1, Figure S3). In inductive link prediction, PPI-GNN and DeepTrio achieve significantly lower performance than UPNA-PPI (see Table 2, Figure 4B). Furthermore, we borrow two performance metrics from recommender systems into PPI prediction to assess the quality of the ranking made by PPI prediction models. We define PPIHits@TopK as the number of true interactions (precision) identified by an ML model in the top K predictions. We also define, PPNIHits@BottomK, which quantifies the number of true non-interactions identified in the bottom K predictions of a model. These two metrics bolster our confidence in the ranking provided by the model, which is associated with the separability of the predictions for the PPI and PPNI in test data. UPNA-PPI establishes itself as an excellent ranking tool for identifying PPI in terms of both metrics (PPIHits@Top1000: 0.92 ± 0.015 and PPNIHits@Bottom1000: 0.69 ± 0.08). In Figure 4C, we observe how DeepTrio creates overlapping predictions for the true interactions and true non-interactions. The incorporation of PPNI at the output of the configuration model combined with pair-wise learning improves the separation between these distributions and helps UPNA-PPI predict the true interactions with higher probabilities. Finally, filtering the PPNI with L3 counts (topological PPNI) separates the UPNA-PPI output distributions, confirming the ability of UPNA-PPI to learn the latent protein-protein interaction patterns driven by various molecular complementarity mechanisms. These observations provide an ablation study for the topological PPNI and showcase the strength of the CL3 approach in creating hard negatives.

**Table 1:** State-of-the-art PPI Prediction Models in Transductive and Inductive Link Prediction. In transductive link prediction, nodes are shared between train and test datasets. State-of-the-art PPI prediction models, PPI-GNN and DeepTrio, achieve excellent transductive link prediction performance. However, in inductive tests, when encountered with never-before-seen proteins, the performances of both models diminish significantly.

| Model | Transductive |  | Inductive |  |
| --- | --- | --- | --- | --- |
|  | AUROC | AUPRC | AUROC | AUPRC |
| PPI-GNN | $0.98 \pm 0.044$ | $0.98 \pm 0.042$ | $0.50 \pm 0.00$ | $0.74 \pm 0.03$ |
| DeepTrio | $0.93 \pm 0.003$ | $0.94 \pm 0.005$ | $0.59 \pm 0.06$ | $0.62 \pm 0.07$ |

**Table 2:** State-of-the-art PPI Prediction Models vs. UPNA-PPI in Inductive Link Prediction. Our proposed pipeline UPNA-PPI combining topological PPNI and unsupervised pre-training on protein representations, significantly improves inductive link prediction performance in PPIs compared to two state-of-the-art PPI prediction models PPI-GNN and DeepTrio.

| Model | AUROC | AUPRC | Hits@Top1000 | Hits@Top10,000 |
| --- | --- | --- | --- | --- |
| UPNA-PPI | $0.79 \pm 0.007$ | $0.87 \pm 0.009$ | $0.92 \pm 0.015$ | $0.87 \pm 0.010$ |
| PPI-GNN | $0.50 \pm 0.0$ | $0.74 \pm 0.03$ | $0.67 \pm 0.07$ | $0.06 \pm 0.006$ |
| DeepTrio | $0.59 \pm 0.06$ | $0.62 \pm 0.07$ | $0.25 \pm 0.008$ | $0.024 \pm 0.001$ |

On a similar token, the recent discovery of shortcut learning [53] has raised many questions about the reliability of ML predictions and inductive tests have gained significant attention in generalizability and interpretability [51, 54]. RAPPPID [60] is a recently proposed PPI prediction model that uses inductive learning to improve the generalizability of PPI prediction. RAPPPID uses sequential neural models (LSTM) [95] to embed the protein amino acid sequences and feed them to a downstream decoder, which is trained for an inductive setting in an end-to-end fashion. Yet, the regularization of the protein embeddings limits the generalizability of RAPPPID within certain protein datasets and protein classes. Performance of RAPPPID significantly reduces in a transfer learning scenario from STRING [13] to BioLip [96]. In Table 3, we see that UPNA-PPI achieves comparable performance to RAPPPID for inductive link prediction within similar protein families, while performing better than RAPPPID in a transfer learning setting from STRING to BioLip. Data-specific regularization hinders RAPPID from achieving superior generalizability across datasets and protein families, which is ameliorated in UPNA-PPI by unregularized and unsupervised protein representations, independent of the training dataset.

**Table 3:** UPNA-PPI vs. RAPPPID in inductive link prediction and transfer learning across protein families and peptides. RAPPPID is an inductive PPI prediction model which uses sequential models and regularization for protein representations. Although, UPNA-PPI and RAPPID achieve comparable performances in inductive link prediction, UPNA-PPI offers stronger prioritization as depicted by Hits@TopK. Furthermore, in the transfer learning setting of training on STRING DB and testing on BioLip, which contains proteins from different families than STRING DB, RAPPPID fails and achieves performance similar to a naive Bayes classifier on unseen data. However, UPNA-PPI can make meaningful PPI predictions in this transfer learning setting, showing improved generalizability across protein families.

| Model | Inductive |  |  |  | Transfer learning<br>AUROC |
| --- | --- | --- | --- | --- | --- |
|  | AUROC | AUPRC | Hits@Top1000 | Hits@Top10,000 |  |
| UPNA-PPI | 0.89 | 0.90 | 0.98 | 0.93 | 0.73 |
| RAPPPID | 0.80 | 0.81 | 0.77 | 0.13 | 0.55 |

In Figure 4D, we plot the UPNA-PPI prediction for all experimentally validated PPIs against the number of L3 paths induced by these pairs in the PPI network. A robust correlation is evident between the UPNA-PPI predictions and the L3 counts, with r_*Spearman*_ = 0.48. This observation confirms that UPNA-PPI effectively captures the complementarity-driven mechanism underlying L3 in PPIs based on protein amino acid sequences. Moreover, as depicted in Figure 4E, the UPNA-PPI predictions for the majority of the experimentally validated PPIs surpass the optimal classification threshold (0.476 ± 0.049).

### Robustness and Sensitivity of UPNA-PPI

The robustness of link prediction with the introduction of noise and adversarial attacks in the network provides insight into the reputability of machine learning models [97, 98]. Robustness of PPIs have been studied from various aspects, including randomized null models [99], temperature dependence of stable protein complexes [100], and the removal of hub proteins [101]. PPI networks are topologically more robust under four common types of perturbations (i) network nodes are randomly removed (failure), (ii) the most connected node is successively removed (attack), (iii) interaction edges are rewired randomly, and (iv) edges are randomly deleted [102]. In Figure 5A, we show how L3 paths (4-cycles) in PPI make the networks robust under degree preserved edge swap [103]. In Figure 5B, we observe that the inductive link prediction performance of UPNA-PPI does not fluctuate under degree-preserved edge swap in train and validation datasets, while keeping the test data unchanged. This form of robustness is inherent to the PPI network topology, and degree-preserved edge swap is insufficient for evaluating the robustness of any ML-based PPI prediction model.

**Figure 5:**
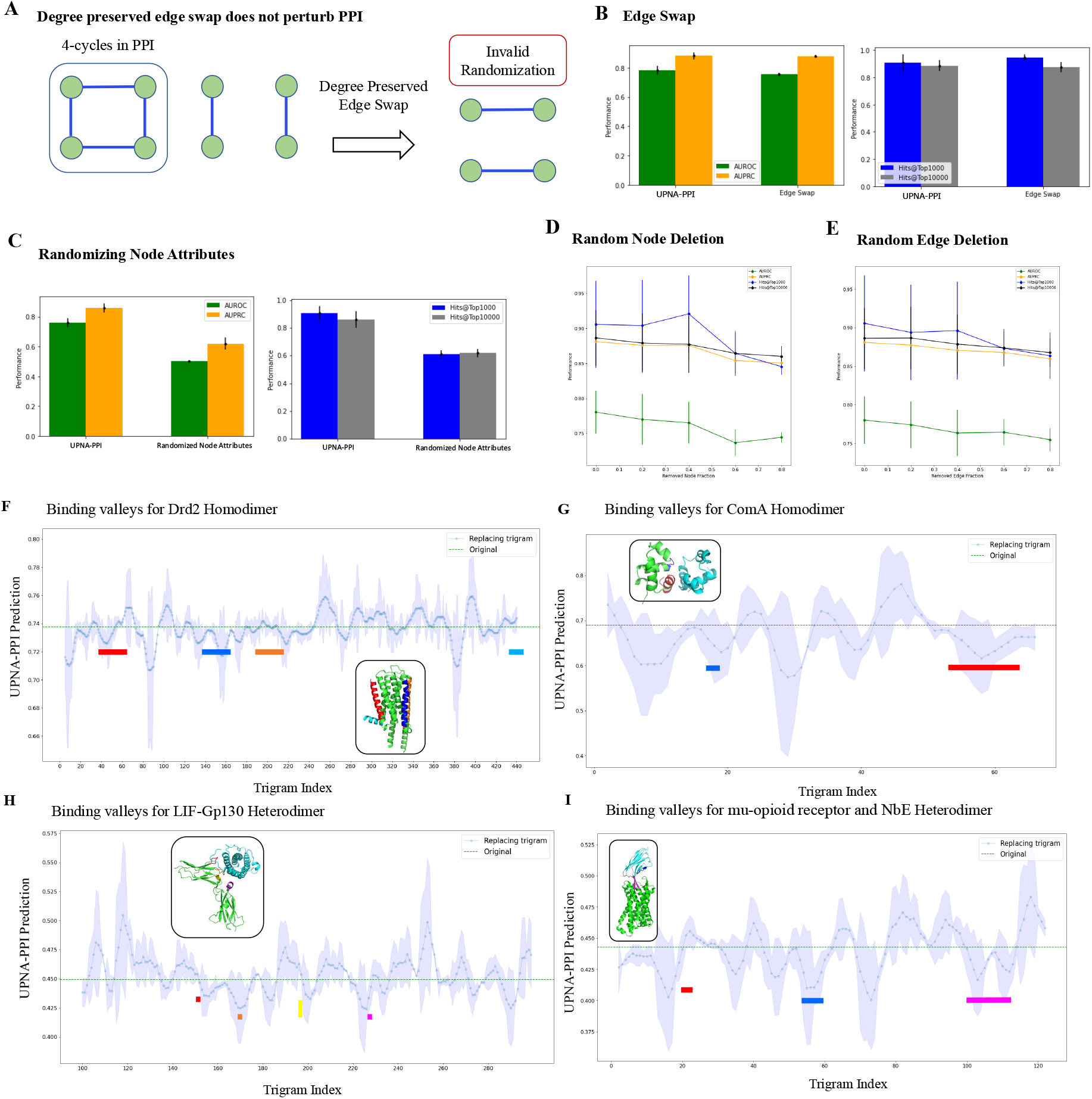
Robustness and Interpretability of UPNA-PPI. **(A)** Degree preserved edge swap is a widely used method to randomize graphs for studying robustness. However, since PPI networks are enriched with 4-cycles, degree preserved randomization frequently duplicates existing interactions in PPI instead of creating random edges. **(B)** In the robustness study we perturb train and validation datasets while keeping the test dataset unchanged. The inductive performance of UPNA-PPI is invariant to degree-preserved randomization. **(C)** We replace ProtVec embeddings with 100-dimensional random vectors where the value for each dimension is randomly selected from a uniform distribution U[0, 1]. We observe inductive performance similar to a naive Bayes classifier under such randomization, confirming that UPNA-PPI learns interactions leveraging the embeddings of the amino acid sequences. **(D)** We randomly remove nodes (proteins) from the training and validation datasets. We do not observe significant fluctuation in UPNA-PPI inductive performance, which confirms the ability of UPNA-PPI to learn from limited data and generalize to new proteins. **(E)** Similar to (D), we randomly remove edges (interactions) from the training and validation datasets. We do not observe significant fluctuation in UPNA-PPI inductive performance, which confirms the ability of UPNA-PPI to learn from limited data and generalize to new proteins. **(F)** We ran an ablation study on the amino acid sequence of DRD2 to identify potential trigrams where interaction takes place for creating Drd2 homodimer. In this ablation study, each amino acid trigram is replaced with the Out-of-Vocabulary embedding from ProtVec, while keeping the other amino acid sequence in the input of UPNA-PPI unchanged. We observe multiple valleys in the binding probability profiles. These valleys correspond to the interfeces TM4/TM5 and TM1/H8 which have been identified experimentally using Cys-crosslinking and FRET. **(G)** We repeat a similar process for another homodimer of transcription factor ComA. **(H)-(I)** We run abltation study on two heterodimers consisting of protein pairs LIF-Gp130 and Mu-opioid receptor-NbE. The interaction locations on protein complexes LIF-Gp130 and mu-Opioid receptor-NbEHH are marked in the figures, which overlap with valleys predicted by UPNA-PPI. In all of the above scenarios, we observe that the valleys with lower standard deviation obtained from 5-folds of UPNA-PPI correspond to the true binding locations. Therefore, the valleys on which 5-fold of UPNA-PPI agree are the binding locations with a higher confidence.

Next, we test the sensitivity of UPNA-PPI on the protein representations. We replace the 100-dimensional ProtVec embeddings with vectors whose entries are drawn at random from a uniform distribution U[0, 1]. In Figure 5C, we observe that UPNA-PPI performance drops significantly when the protein representations are randomized and the performance is comparable to a naive Bayes’ classifier. This confirms that UPNA-PPI learns PPI mechanisms leveraging the protein amino acid sequences and does not refuge to any form of shortcut learning.

Finally, we test the robustness of UPNA-PPI under random node and edge deletion. We perturb the train and validation datasets while keeping the test data unchanged. In Figures 5D and 5E, we observe slight decay in the inductive test performance of UPNA-PPI under random deletion of nodes and edges from the training and validation PPI. However, even after deleting 80% of training nodes and edges, UPNA-PPI is still able to achieve commendable inductive link prediction performance. This confirms the high generalizability of the proposed ML pipeline under data scarcity and the power of topological negatives in capturing the complementary mechanisms behind protein interactions.

#### Interpretability and Identifying Interaction Locations

Finally, the molecular interpretability of PPIs is crucial for understanding their biological significance, their role in various cellular processes, and their potential as drug targets. We developed an ML-based approach for identifying putative interaction regions in a given pair of putative interacting proteins by ablating their sequences [104]. Keeping the amino acid sequence of one protein fixed at a time, we mutate each amino acid trigram of the other protein by replacing it with the Out-of-Vocabulary (OOV) probing with in-vocabulary example (IVE) entry in ProtVec, and observe the change in UPNA-PPI output prediction probability. Note that, the OOV embedding vector in ProtVec is obtained by the average of all the trigram embedding vectors [85]. The valleys of the binding probability profiles correspond to the potential binding sites on the protein sequence. We validate this method on experimentally validated self-interacting protein pairs Drd2 [105] and ComA [106] heterodimeric complexes of proteins: Leukemia inhibitory factor (LIF) in complex with gp130, and mu-opioid receptor bound to NbE. In Figure 5 (F-I), we mark the valleys identified by UPNA-PPI which correspond to the binding interfaces identified by experimentally for these complexes. For Drd2 the interacting TM helices have been reported as self-interactions between a region around TM4, and another one in TM1 utilizing Cys-crosslinking and FRET [105] (Figure 5F). The transcription factor Competence protein A (ComA) dimerization has been determined by NMR showing self-interaction mostly on C-terminal helix alpha-10 [106]. LIF interaction interface with gp130 was determined by X-ray crystallography [107]. Finally, the binding site of NbE on the orthosteric site of the mu-opioid receptor was also determined with X-ray crystallography [108].

#### Novel interactions for G protein-coupled receptors

Physiologically and pharmacologically relevant G-protein-coupled receptors (GPCRs) are the target of several marketed drugs [109]. They mediate physiological responses to hormones, neurotransmitters, and environmental stimuli. Recent high-resolution structural studies have shed light on the molecular mechanisms of GPCR activation and constitutive activity. Over the past three decades, significant progress has been made in understanding GPCRs, from pharmacology to in vivo function. However, only G protein-coupled receptor-G protein interactions have been explored extensively [110]. We used UPNA-PPI to predict potential homodimers and heterodimers between GPCRs. We consider 170 family A GPCR proteins for this prediction task. In Figure 6, we visualize the GPCR interaction network from top 100 UPNA-PPI predictions. The network consists of 28 GPCR protein nodes with 8 homodimers and 92 heterodimers. Furthermore, since UPNA-PPI is not trained on self-interactions, we evaluated the homodimeric predicted by UPNA-PPI with AlphaFold-Multimer predicted homodimer structures [111, 112]. In Table 4, we summarize the AlphaFold average of predicted local distance difference test (pLDDT) score, atoms at interface, and interface pLDDT score for top and bottom homodimer predictions made by UPNA-PPI (see Figure 6). Segregation of the potential GPCR interactions predicted by UPNA-PPI is in agreement with more computationally expensive results from AlphaFold-Multimer. Hence, UPNA-PPI can be used as a potential high throughput tool for novel PPI prediction.

**Table 4:**
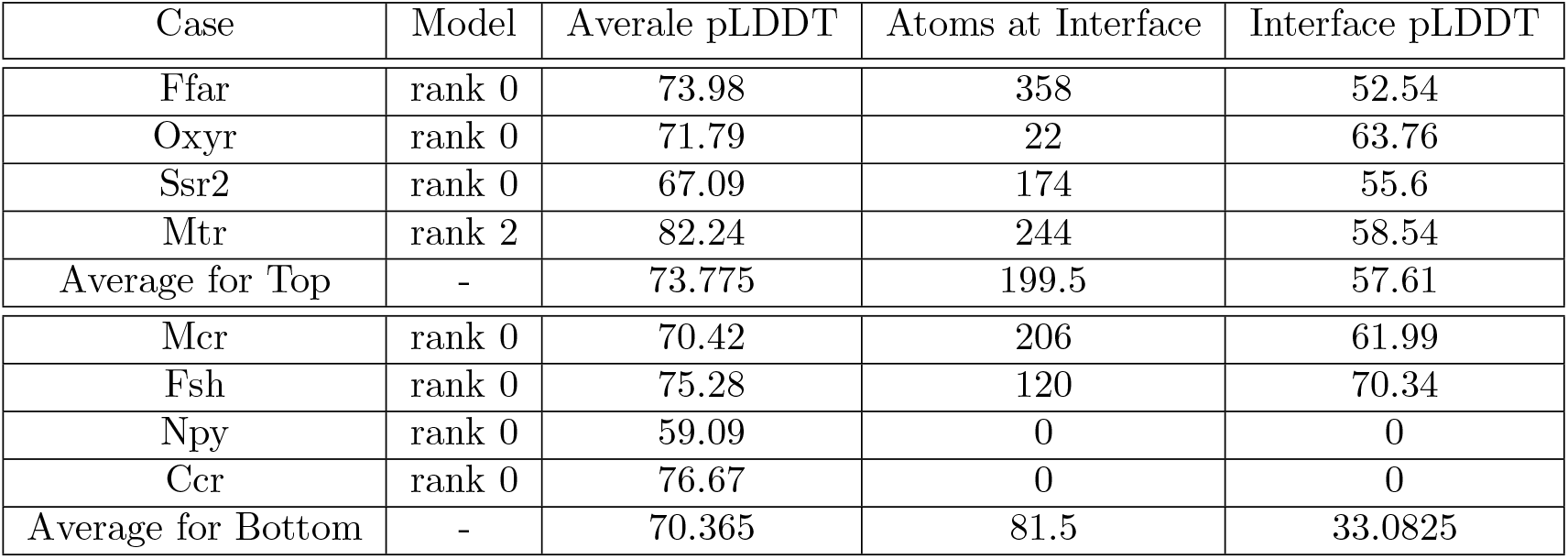
UPNA-PPI validation with AlphaFold Multimer. While UPNA-PPI is trained on heterodimers, we test the transferability of UPNA-PPI by predicting homodimers. For this experiment, we use understudied GPCR proteins. The top GPCR homodimer predictions from UPNA-PPI include the protein Fafr, Oxyr, Ssr2, and Mtr. The bottom predictions for self-interaction prediction include the proteins Mcr, Fsh, Npy, and Ccr. We run AlphaFold-Multimer simulations for the listed proteins and summarize the observations in the table below. We observe that, on average, the top self-interaction predictions from UPNA-PPI show more atoms at the interaction interface and higher pLDDT (confidence) at the interface compared to the bottom predictions. Furthermore, while the average pLDDT values are comparable between top and bottom predictions, interface pLDDT is higher for the top predictions, suggesting that UPNA-PPI is able to learn the interfaces of interaction leveraging only the amino acid sequences.

| Case | Model | Average pLDDT | Atoms at Interface | Interface pLDDT |
| --- | --- | --- | --- | --- |
| Ffar | rank 0 | 73.98 | 358 | 52.54 |
| Oxyr | rank 0 | 71.79 | 22 | 63.76 |
| Ssr2 | rank 0 | 67.09 | 174 | 55.6 |
| Mtr | rank 2 | 82.24 | 244 | 58.54 |
| Average for Top | - | 73.775 | 199.5 | 57.61 |
| Mcr | rank 0 | 70.42 | 206 | 61.99 |
| Fsh | rank 0 | 75.28 | 120 | 70.34 |
| Npy | rank 0 | 59.09 | 0 | 0 |
| Ccr | rank 0 | 76.67 | 0 | 0 |
| Average for Bottom | - | 70.365 | 81.5 | 33.0825 |

**Figure 6:**
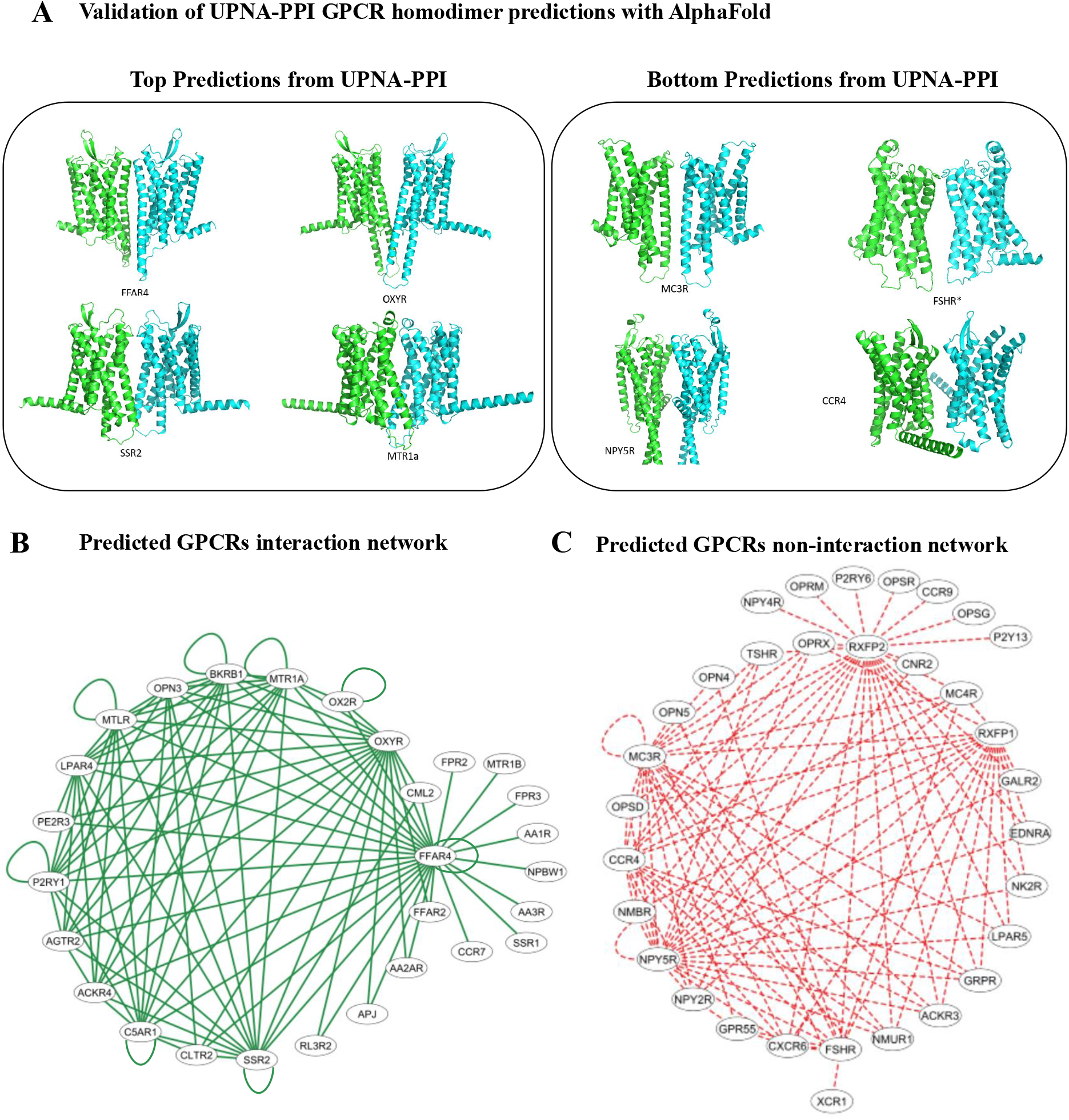
UPNA-PPI Predictions for G-protein coupled receptors (GPCRs). **(A)** We validate top and bottom self-interaction predictions for GPCRs from UPNA-PPI with AlphaFold multimer. We observe that AlphaFold predicts a higher number of atoms at the interaction surface and pLDDT at the surface for the top predictions compared to the bottom predictions (see Table 4), which validates the agreement between UPNA-PPI and AlphaFold. **(B)** Interaction network of GPCRs. From top 100 UPNA-PPI predictions we construct the interaction network of GPCRs. We predict 8 homodimers and 92 heterodimers involving 28 GPCR proteins. **(C)** Similarly, we construct a non-interaction network with bottom 100 predictions from UPNA-PPI.

## DISCUSSION

While machine learning has widely been used in protein-protein interaction prediction, generalizing these models to novel proteins, transferring across protein families, and interpretability of predictions from a molecular standpoint have emerged as the major pitfalls of these models. The scarcity of high-quality, biologically relevant hard PPNI sampling has been a major hindrance for machine learning models to learn interaction mechanisms from protein structural patterns. We have proposed a novel approach to leveraging the PPI network topology in sampling PPNI contingent on the complementarity-driven generation mechanisms of protein-protein interactions. We interpret topological PPNI geometrically by leveraging the hyperbolic space and make a connection between the complementarity of PPI and the evolution of proteins. Our PPNI sampling approach combined with unsupervised pre-training of protein representation not only improves the generalizability of PPI prediction but also improves the transferability of machine learning prediction across protein families. UPNA-PPI is also able to identify potential pocket locations on the amino acid sequences, bolstering the molecular interpretability of machine learning prediction. Furthermore, the robustness of UPNA-PPI under random node and edge removal strengthens the notion of generalizability under data scarcity [113].

Complementarity-driven networks induce even-length cycles [42]. We have developed a novel negative sampling strategy leveraging the 4-cycles in PPIs. This approach can be extended to other complementarity-driven networks by exploring the even-length cycles enriched in the networks. Furthermore, there has been much research in identifying and counting 4-cycles in directed graphs [114,115] and sparse graphs [116]. Integrating these algorithms to the topological PPNI methodology can ameliorate the need for the configuration model and help us derive more hard PPNI samples. We also hypothesize that PPI networks from different species should be treated individually to create topological PPNI. Considering the combined PPI of humans and other species would hinder us from capturing meaningful evolutionary patterns and the complementary nature of the network.

In its exploration of drug-target interaction networks, AI-Bind [54] employed network science to comprehend topological shortcuts and generate negative samples. In contrast, UPNA-PPI extends the application of network science to ML models operating on protein-protein interaction networks. UPNA-PPI introduces a novel methodology for negative sampling by leveraging higher-order network properties. Beyond supplying valuable hard negatives applicable to a diverse array of ML models, UPNA-PPI pioneers a new research direction that advocates the use of network topology in negative sampling. This innovative approach holds promise for enhancing the generalizability, robustness, and interpretability of graph machine learning methodologies on a wide range of networks.

## METHODS

### Merging and Consolidating PPI Databases

Consolidating the human interactome using the existing experimental databases is challenging [117]. While the majority of the existing PPI databases often report only the gene identifiers (gene symbols [118], Entrez ID [119], Ensembl ID [120], etc.), newer databases like BioPlex [12] report the protein IDS (UniProt ID [25], PDB ID [121], etc). Ambiguity in gene-to-protein mapping combined with the annotation inconsistency in these databases makes the task of merging the PPI data difficult [122]. Furthermore, the majority of the computational tools for exploratory biology, including network science-inspired disease module identification algorithms [123, 124], demand a coarse-grained representation of the interactome. ML-based PPI prediction methods [56, 60, 94] use UniProt IDs to specify proteins, and hence disregard a big portion of available experimental PPI data. For the aforementioned reasons, we coarse-grain human protein IDs from UniProt to gene Entrez IDs. Establishing a one-to-one correspondence between human genes and amino acid sequences involves selecting the longest amino acid sequence from all protein isoforms associated with a given gene. We have merged multiple experimental PPI databases such as BioPlex [12], STRING [13], APID [14], BioGRID [15], CoFrac [16], CORUM [17], HuRI [18], HINT [19], HIPPIE [20], and BKB [125] to obtain our interaction data.

### GoldPPI

To accumulate high-confidence PPI data, we filter the experimental PPI databases for samples with multiple experimental validation. We use three AP-MS databases, which report the number of experiments/publications validating the interaction: APID [14], HINT [19], and HIPPIE [20]. Thereafter, we filter for the PPI samples which have been observed in at least 3 experiments. Thus, we obtain the high confidence PPI, which we use in UPNA-PPI validation and test datasets.

### Negatome PPNI

The majority of the PPI databases do not report the non-interactions. Hence, there is a lack of PPNI data in the literature, which has led the majority of the ML approaches to use random negative sampling or subcellular compartmental negatives. We use high-confidence PPNI from Negatome 2.0 [126] and NVDT [26]. These samples have been used in the validation and test of UPNA-PPI.

### Traditional/Simple Unipartite Configuration Model

The SCM is an exponential random graph model with the probability of observing a graph

Configuration 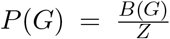, where 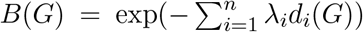, the Lagrange multipliers {λ_*i*_} are such that 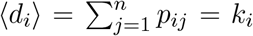 for all nodes *i* ∈ {1, 2, …, *n*}, the Boltzmann factor 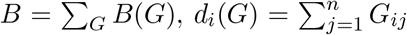 is the degree of the node *i*, and 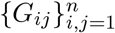 is the adjacency matrix of the training graph *G*. By entropy maximization, we get the link probability between the nodes *i* and *j* as:

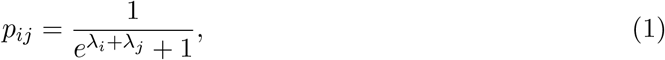

## Data availability

QIAGEN BKB (Biomedical Knowledge Base) PPI data is available upon request to QIAGEN. The UPNA-PPI predictions will be shared upon reasonable request. The PPIs obtained from WikiPathways, STRING, and BioLip is available on Zenodo.

## Code availability

The codes that support the findings of this study are openly available on our GitHub at https://github.com/alxndgb/UPNA-PPI.

## Acknowledgements

We thank the Bioinformatics and Data Science group in Alexion AstraZeneca Rare Disease for funding this project. We are grateful to AstraZeneca’s Scientific Computing Platform (SCP) for providing the computing resources. We thank István Kovács, Assistant Professor in the department of Physics and Astronomy at Northwestern University for fruitful discussions on the structural characteristics of PPIs and the extent of incompleteness in the human interactome. We thank Dr. Guillermo del Angel, the Rare Disease TA head at Center for Genomics Research in AstraZeneca for his constructive feedback and highlighting the importance of negative samples in modeling PPI networks. Lastly, We are grateful to Dr. Gyan Srivastava, Associate Director of Alexion Bioinformatics and Data Sciences, and Andrew Proffitt, Research Scientist at Alexion Rare Disease New Target Discovery and Validation group, for their fruitful discussions and guidance.

## Author Contributions

A.C. and B.R. conceived and designed the project, wrote the manuscript, conducted data curation and preparation, generated the predictions for the network configuration model, performed experiments to identify networked-derived negatives, implemented negative sample generation, developed and testing of UPNA-PPI, assessed the robustness of UPNA-PPI, quantifying the generalizability of PPI predictions, implementation of the interpretability component to predict the active binding locations, and comparisons with SOTA models. P.H. offered biological insights for practical implementations and generalizability of UPNA. N.H.P. prepared the dataset for interpretability analysis by mapping amino-acid sequences of protein pairs to interaction sites and provided biological insights. M.A. and W.R.M. interpreted the results and offered biological insights. P.R. provided insights into subcellular and random negatives and offered biological insights. Y.L. provided IT support. W.D. interpreted the results and offered biological insights. J.C.M. designed the study on GPCRs, executed AlphaFold validations, conducted computational structural simulations for the interpretability of UPNA-PPI predictions, and provided chemical and biological insights. T.E.R. provided guidance on designing the machine-learning experiments, interpreting CL3 hypothesis through the lens of complementarity in complex networks, and writing the manuscript.

## Supplementary Information

### 1 Revisiting Topological Shortcuts in PPI Network

Most real-world networks, such as Protein-Protein Interaction (PPI) [1] and Drug-Target Interaction (DTI) [2] networks, typically exhibit a degree distribution that follows a heavy-tailed distribution [3]. This means that the majority of nodes in these networks have only a small number of links (interactions or samples) in the training datasets, while a small number of hub nodes have a large number of connections. During the training phase of a link prediction model, the model tends to focus more on these hub nodes than on the nodes with lower degrees. In traditional machine learning scenarios, train and test datasets are created by randomly dividing the edges within the network [4]. As a result, the majority of edges in both the training and test datasets involve these hub nodes. Machine learning models become proficient at learning the neighborhood characteristics of these hub nodes in terms of their degree information [5,6]. Consequently, they can make accurate predictions for the edges associated with these hubs, leading to excellent test performance. Figure S2 demonstrates the need for the inductive scenario in link prediction tasks. The transductive scenario, most frequently used to evaluate the performance of AI models, favors learning topological shortcuts instead of the Mechanisms of Actions (MoA) behind the emergence of structural topology in networked data (Figure S2B). In other words, state-of-the-art (SOTA) models leverage the neighborhood topology of a node (protein or drug) to make new interaction predictions, which are often truant of biological and molecular interpretability. However, in an inductive setting (see Figure S2A), the SOTA models struggle when dealing with the low-degree nodes, since they are forced to leverage the molecular patterns for learning interactions. Hence, hubs play the main role in the misleading high performance of the SOTA models. However, in Erdős-Rényi (ER) graphs [7], there are no hubs, and all nodes are indistinguishable in terms of their neighborhood degrees (see Figure S2B). Thus, link prediction models achieve performance similar to a naive Bayes classifier (AUROC ≈ 0.5) when trained and tested on ER graphs (see SI Section 1 of [5]). Furthermore, AI models trained under transductive scenarios are unable to perform well even in transductive link prediction tasks for low-degree nodes [5]. This highlights the models’ limitations in extracting meaningful patterns from the molecular structures of proteins, resulting in poor inductive performance, revealing the misleading nature of transductive tests, and insufficient biological interpretability of the machine learning predictions. Likewise, current machine learning models demonstrate outstanding effectiveness when analyzing common diseases that benefit from abundant data available in established databases [8]. Nonetheless, when it comes to rare diseases [9], the limited data availability restricts the utility of these models in making valuable predictions (see Figure 2C).

#### 1.1 State-of-the-art PPI Prediction Models in Transductive Tests

Protein-protein interactions (PPIs) exhibit a complex hierarchy of generation mechanisms. At the top level, we have Preferential Attachment (PA) [3], a concept that plays a crucial role in a link prediction model’s ability to excel when it comes to predicting connections involving hub proteins. However, it is important to note that PA itself arises as a consequence of certain proteins having a greater number of binding pockets and findings in experimental studies. SOTA models, despite their sophistication, often fall short of understanding the true underlying mechanisms by leveraging the molecular structures of proteins. It’s during inductive tests that these models are compelled to uncover this genuine generative process, which, in turn, enhances the model’s interpretability. This assertion is substantiated through an experiment involving the introduction of random amino acid sequences onto a DTI network, where the state-of-the-art binding prediction model DeepPurpose [10] model achieves exceptional transductive performance, showcasing the capacity to discern the authentic generation mechanism.

In Figure S3B we visualize the heavy-tailed degree distribution of the QIAGEN BKB PPI [11]. We consider two state-of-the-art PPI prediction models, DeepTrio [12] and PPI-GNN [13]. We develop a unipartite duplex configuration model (see Section S2) [14], which takes as input only the degree sequences from the protein-protein interaction and non-interactions (random negative sampling) used in the training of DeepTrio and PPI-GNN (see Figure S3B). We observe that using only the degree information of the proteins and being completely blind to the protein structures (amino acid sequences), the configuration model achieves comparable performance to the SOTA models (Figure S3C). This experiment confirms that the SOTA models are unable to learn from the protein amino acid sequences and resort to topological shortcuts [5], hence lack biological interpretability in the predictions.

#### 1.2 Sealing Information Leakage, Inductive Tests, and Generalizability

Following the common formulation of GraIL [15], Park et al. [16], and Chatterjee et al. [6,17], we evaluate the performance of the SOTA models in different test scenarios. We have already seen that in the transductive setting, proteins are shared between train and test datasets in a transductive setting (Figure S2A). In a semi-inductive setting, one protein of the test edge is present in train data, and the other one is unseen during training. In an inductive link prediction setting, both proteins of the test edge are absent in training. In Table 1 in the main text, we observe a gradual lowering of the test performance of the SOTA models as we move from transductive to inductive tests.

This observation is primarily driven by the reduction of topological information leakage between train and test data. To develop a generalizable AI model, learning PPI from the protein structures instead of leveraging the topology that inflates the performance, we need to mitigate shortcut learning and make protein representation learning independent of the PPI training data. Furthermore, regularization has been proven to be an important tool for improving generalizability in computer vision [18]. Yet, regularization is based on the assumption that similarity between a pair of entities implies interaction, an assumption that is true in image processing and social sciences [19], yet similar molecules do not necessarily always interact in PPI and other biological systems [20]. Although [19] is applicable for identifying similarity between images, the similarity of proteins does not imply their interaction, and hence the regularization-based approach to overcome negative sampling is not applicable in PPI prediction. Thus, we leverage the network topology of the PPI network to create high-quality PPNI for training our AI pipeline.

### 2 Unipartite Duplex Configuration Model

#### 2.1 Overview

Protein-protein annotations naturally form an integral part of a unipartite duplex network, which can be conceptualized as a network comprising a set of nodes representing all proteins. This network operates on two distinct layers, with each layer corresponding to a specific type of interaction occurring between the same pair of nodes, as described in Menichetti et al. [14]. Specifically, Layer 1 characterizes positive interactions, while Layer 2 records negative interactions (see Figure S3B).

The multilink notation, denoted as **m**, is employed to encode the pattern of links connecting two nodes across different layers. Notably, **m** = (1, 0) signifies positive interactions, **m** = (0, 1) represents negative interactions, **m** = (0, 0) signifies the absence of any interaction, and **m** = (1, 1) is mathematically prohibited, as it is impossible for both positive and negative interactions to coexist for the same pair of proteins.

In our analysis, we utilize the canonical unipartite duplex null model, which ensures the conservation, on average, of the number of positive and negative annotations associated with each node. This model also appropriately rewires positive and negative links while avoiding the occurrence of forbidden configurations. By applying entropy maximization with constraints, we derive analytical expressions for the probability of each multilink configuration and the conditional probability of observing a positive binding event once an annotation is reported.

#### 2.2 Mathematical Formulation

Let 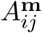 denote the multi-adjacency matrix representing the unipartite duplex network of proteins {*i*} and {*j*}, with elements equal to 1 indicating the presence of a multilink **m** between proteins *i* and *j*, and 0 otherwise. We define the multidegree of protein *i* as:

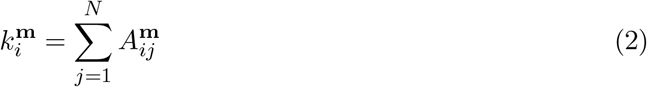

Here, *N* represents the total number of proteins.

A unipartite duplex network ensemble encompasses all duplexes that adhere to specific constraints, such as the expected multidegree sequences defined in Equation (S2). The probability of observing a unipartite duplex network, denoted as 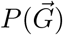, is determined using entropy maximization with multidegree constraints 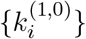 and 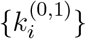, along with their corresponding Lagrangian multipliers 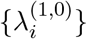 and 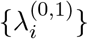, as outlined in [14, 21]. The probability 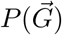 factorizes as follows:

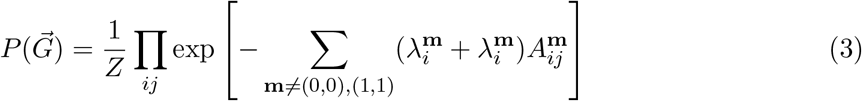

Where:

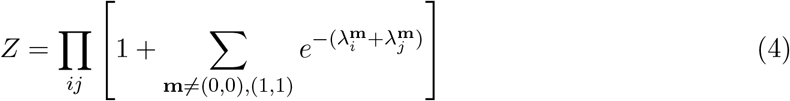

The multilink probabilities 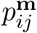 are determined by the derivatives of the logarithm of *Z* with respect to 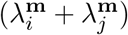 For instance, the probability of observing a positive annotation is expressed as:

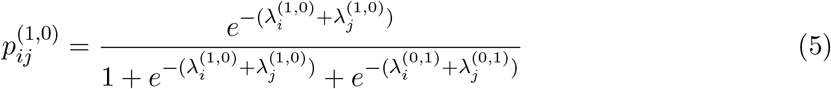

Similarly, the probability of observing a negative annotation is given by:

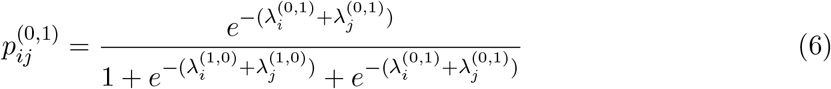

It is important to note that 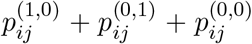 equals 1, indicating a normalization of probabilities.

In this theoretical framework, binding prediction is inherently conditional, focusing on the presence of positive and negative annotations for proteins *i* and *j*. Therefore, 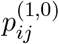 and 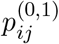 are normalized by the probability of observing a generic annotation, i.e.,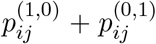. For unseen edges, binding prediction is determined by:

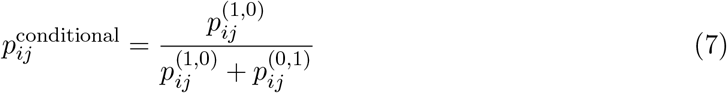

In the case of an unseen protein *j*^*∗*^, the binding probability toward a known compound *i* is calculated as follows:

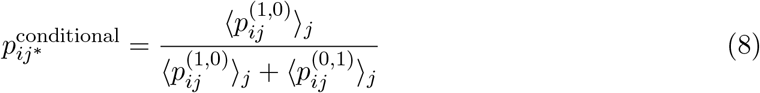

Where ⟨·⟩_*j*_ represents the average over all known proteins.

For unseen proteins *i*^*∗*^ and *j*^*∗*^, the binding probability is determined based on the overall number of positive (*L*^(1,0)^) and negative (*L*^(0,1)^) annotations:

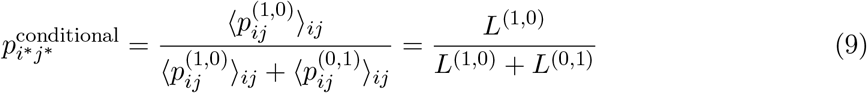

Here, ⟨·⟩_*ij*_ indicates the average over all known protein pairs.

### 3 Annotation Imbalance

The absence of negative samples in PPI databases results in a node-wise class imbalance, where only positive examples are available for the majority of proteins during training. This limitation leads to overpredictions of interactions for these proteins, as the ML models lack exposure to both positive and negative examples [5]. Annotation imbalance is a pervasive issue in interaction databases, as discussed in the context of drug-target binding databases in AI-Bind [5].

To address the annotation imbalance in protein-protein interactions, we utilize topological PPNI. As depicted in Figure S4A, topological PPNI introduces a significant number of negative samples (3,063,605) compared to subcellular compartmental negatives (738,551) and Negatome (2,211). This approach effectively tackles the challenge of annotation imbalance by providing both positive and negative samples for the majority of the proteins in the training, validation, and test sets of UPNA-PPI (see Figure S4B).

### 4. Coefficient of Variation for GPCR Predictions

To further investigate UPNA-PPI output, we looked into the coefficient of variation (for the 5-folds) as illustrated in Figure S5. Coefficient of variation is able to capture the combined effect of the standard deviation and mean of the predictions from 5-folds of UPNA-PPI.

### 5. Alternate protein Embeddings

We evaluate UPNA-PPI against another alternative protein representation learning methodology. Variational autoencoders (VAE) [22] are utilized to derive protein embeddings from amino acid sequences by minimizing both reconstruction loss and distributional loss. We implement VAE-based embeddings following the methodology proposed by Hawkins-Hooker et al. [23]. Initially, we generate one-hot encodings for the amino acid residues and employ these encodings to represent the sequences. Subsequently, a VAE is trained to produce 100-dimensional representations of the sequences. Alternatively, we employ a message-passing neural network (MPNN) [24] on 3D protein structures sourced from the Protein Data Bank (PDB) and those predicted by AlphaFold [25]. However, our examination reveals that Protvec outperforms both VAE and MPNN in inductive testing scenarios (refer to Table S1). Furthermore, we create a simple embedding of the 3D protein structures from the PDB files. We one hot encode the atoms and flatten out the information in the PDB files (atoms, coordinates, and electron densities). We use zero-padding at the end for making the embeddings of same length for different proteins. We observe that these simpler 3D embeddings, although performs poorly compared to Protvec, show improved performance compared to the MPNN approach.

**Table S1:** Comparing Different Protein Embeddings Methods in Inductive Tests. We compare Protvec-based protein representations with VAE and MPNN-based approaches. In both cases, Protvec performs better. Interestingly, a simpler one-hot encoding-based approach on the PDB files provides better performance compared to the MPNN-based protein embeddings.

| Model | AUROC | AUPRC | Hits@Top1000 | Hits@Top10,000 |
| --- | --- | --- | --- | --- |
| Protvec | $0.79 \pm 0.007$ | $0.87 \pm 0.009$ | $0.92 \pm 0.015$ | $0.87 \pm 0.010$ |
| VAE | $0.64 \pm 0.05$ | $0.63 \pm 0.05$ | $0.74 \pm 0.13$ | $0.64 \pm 0.05$ |
| MPNN | $0.59 \pm 0.06$ | $0.62 \pm 0.07$ | $0.25 \pm 0.008$ | $0.024 \pm 0.001$ |
| PDB | $0.74 \pm 0.03$ | $0.72 \pm 0.03$ | $0.68 \pm 0.01$ | $0.55 \pm 0.02$ |

**Figure S1:**
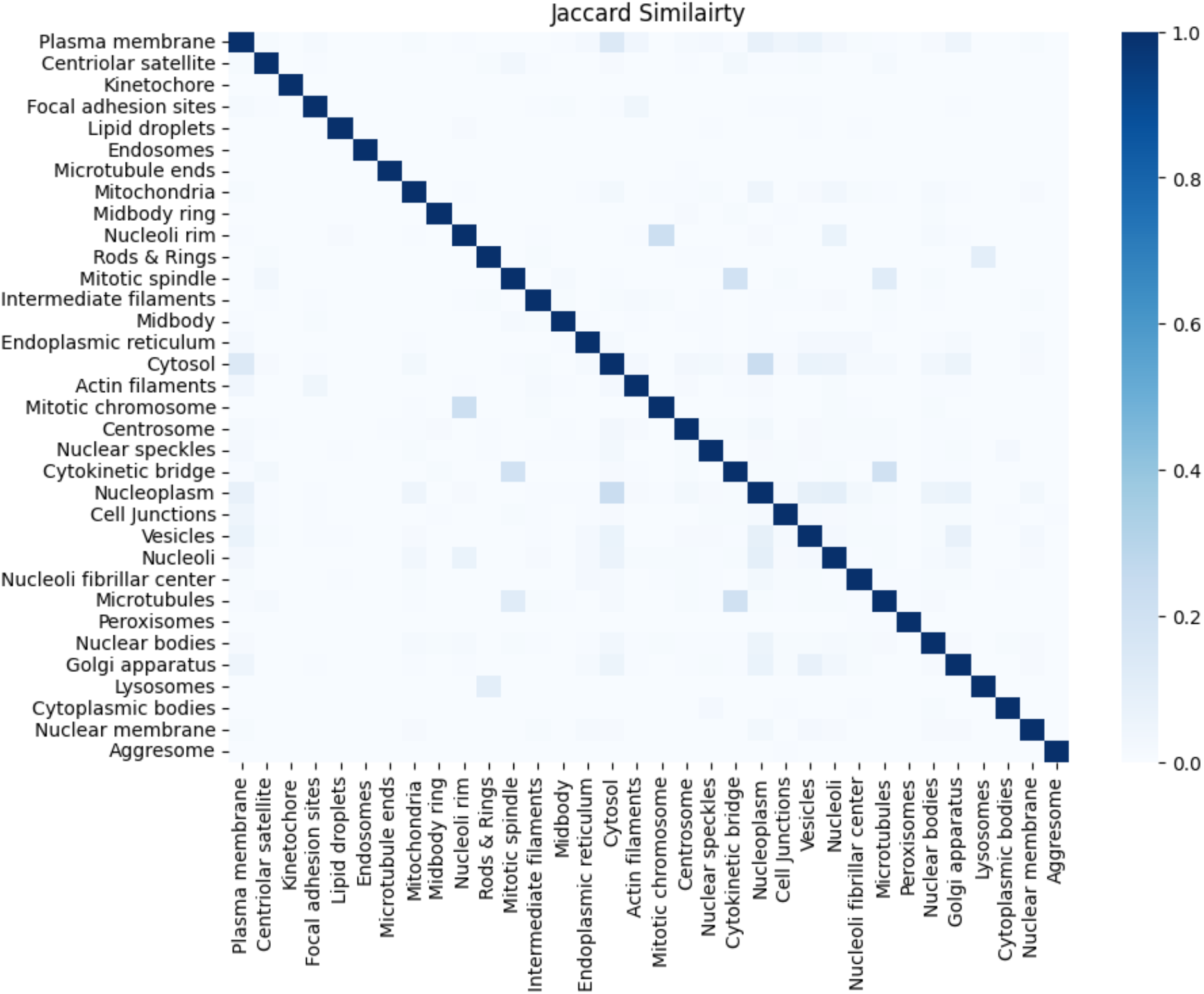
Subcellular Compartmental Negatives (SCN). Protein overlap among 34 subcellular compartments in terms of Jaccard similarity [26]. PPNI pairs are samples from a pair of compartments that have no overlapping protein in the heatmap.

**Figure S2:**
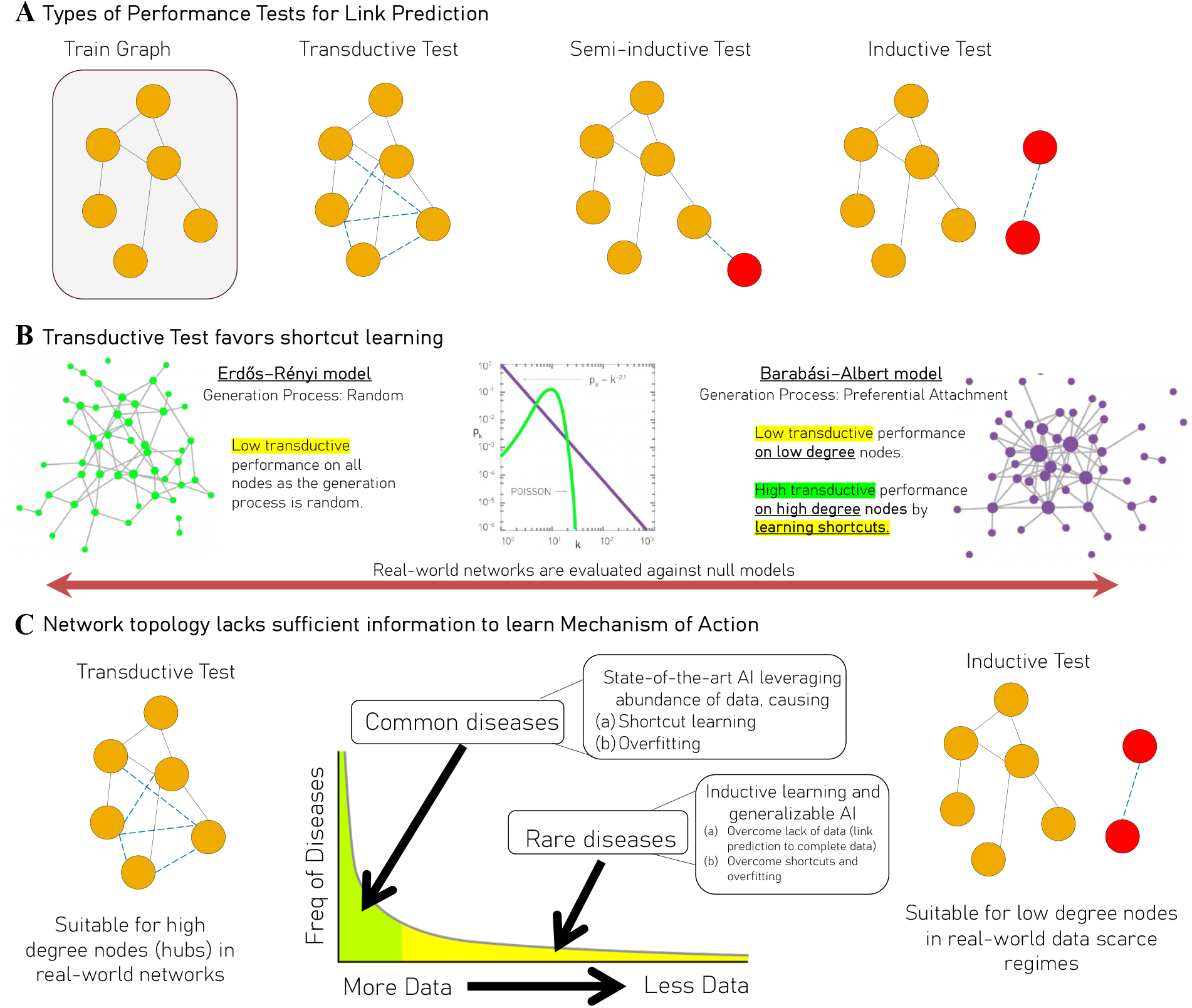
Transductive Test Favors Shortcut Learning. The selection of test scenario can significantly affect the patterns that ML models learn from data. Intuitively, we expect deep models to learn the Mechanisms of Actions (MoA) behind the generation of data, yet it is common that ML models learn shortcuts instead. **(a)** In transductive scenario both the topological structure (patterns of connections) and node attributes are included in the training phase. The semi-inductive scenario partially hides the topological structure from one side of link prediction, by predicting interaction between a node never-seen-before (annotated with red color) and another node that has been seen during the training phase. Lastly, the inductive test prevents the leakage of topological information from train to test/validation sets, conducting link prediction strictly on data-never-seen before, which enforces an AI to learn MoA from node attributes rather than patterns of connections between nodes (shortcuts). **(b)** Real-world networks are often evaluated against two extremes, one is Erdős–Rényi (ER) model where the graph is generated randomly, and the other is Barabási–Albert (BA) model where the graph is generated by enforcing preferential attachment based on the degree of nodes on each iteration of BA model. In ER, the transductive test performs poorly as the graph is generated purely by randomness. In BA model the transductive test only performs well on high degree needs as the classifier learns high degree nodes have a higher chance of having connections. Yet the classifier does not perform well on low degree nodes since the network topology alone does not have enough information to learn the MoA (preferential attachment), which in this case is the parameters used to generate a BA model. **(c)** Transductive scenario may offer insightful predictions when abundant of data is available by facilitating learning the structural topology. Yet, in the data scarce regime, like rare disease area, the performance of transductive scenario is low due to lack of topological structure. In the data scarce regime, learning from node attributes is essential for link prediction, hence we need to enforce learning the MoA behind emergence of structural topology that is domain specific (e.g., social and biological interactions are driven through different MoAs).

**Figure S3:**
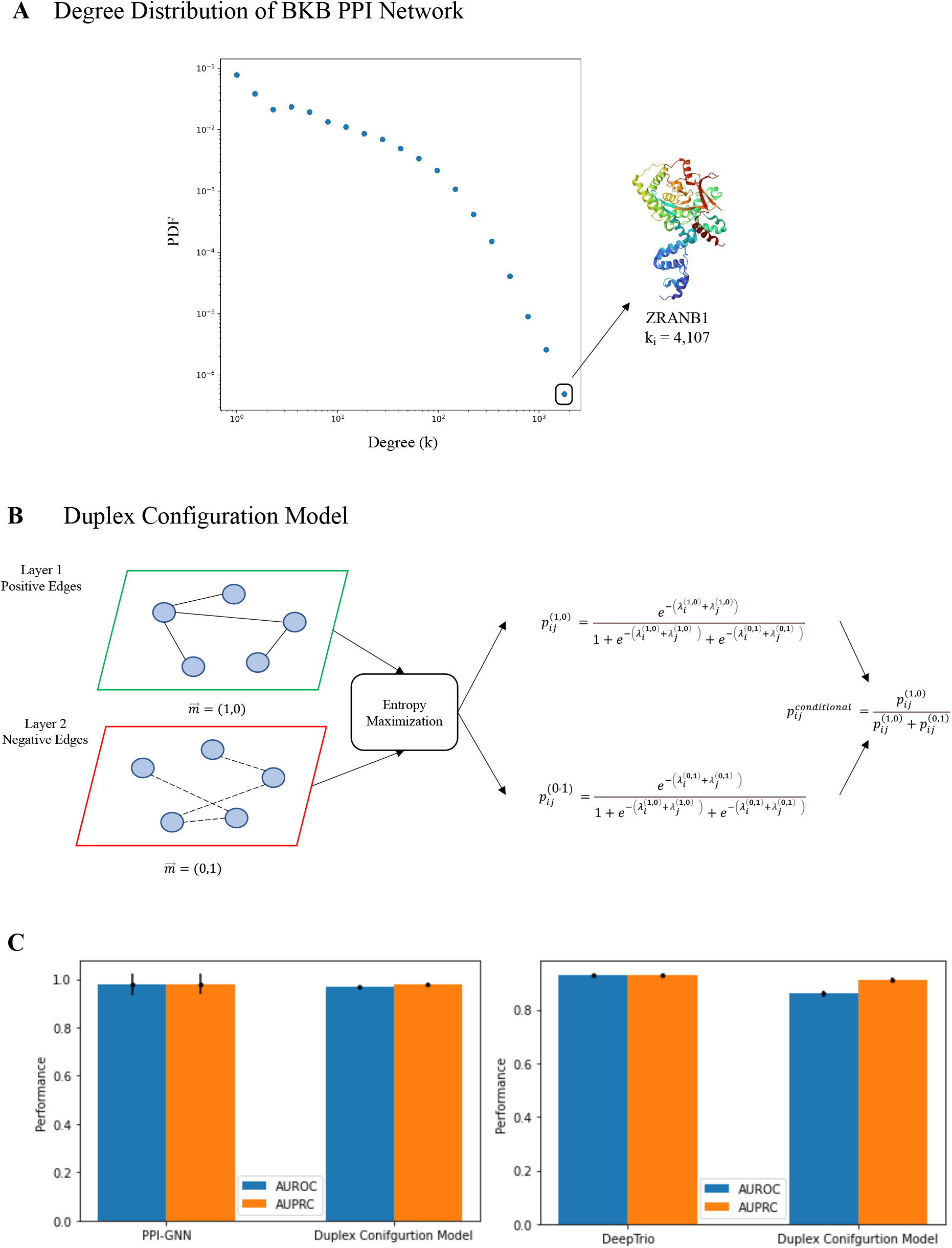
State-of-the-art and Duplex Configuration Model. **(A)** We observe a heavy-tailed degree distribution in the QIAGEN BKB [11] PPI network, which confirms the existence of hubs (high-degree nodes) in the PPI network. For example, ZRANB1 interacts with 4,107 other genes, constituting a hub. **(B)** The unipartite duplex configuration model takes as input PPI training data and random negative samples. By an entropy maximization, we first predict the link probabilities in each layer, which we then combine to obtain the output probability *p*^*conditional*^. **(C)** The duplex configuration model achieves performance comparable to two state-of-the-art PPI prediction models PPI-GNN and DeepTrio in the transductive setting.

**Figure S4:**
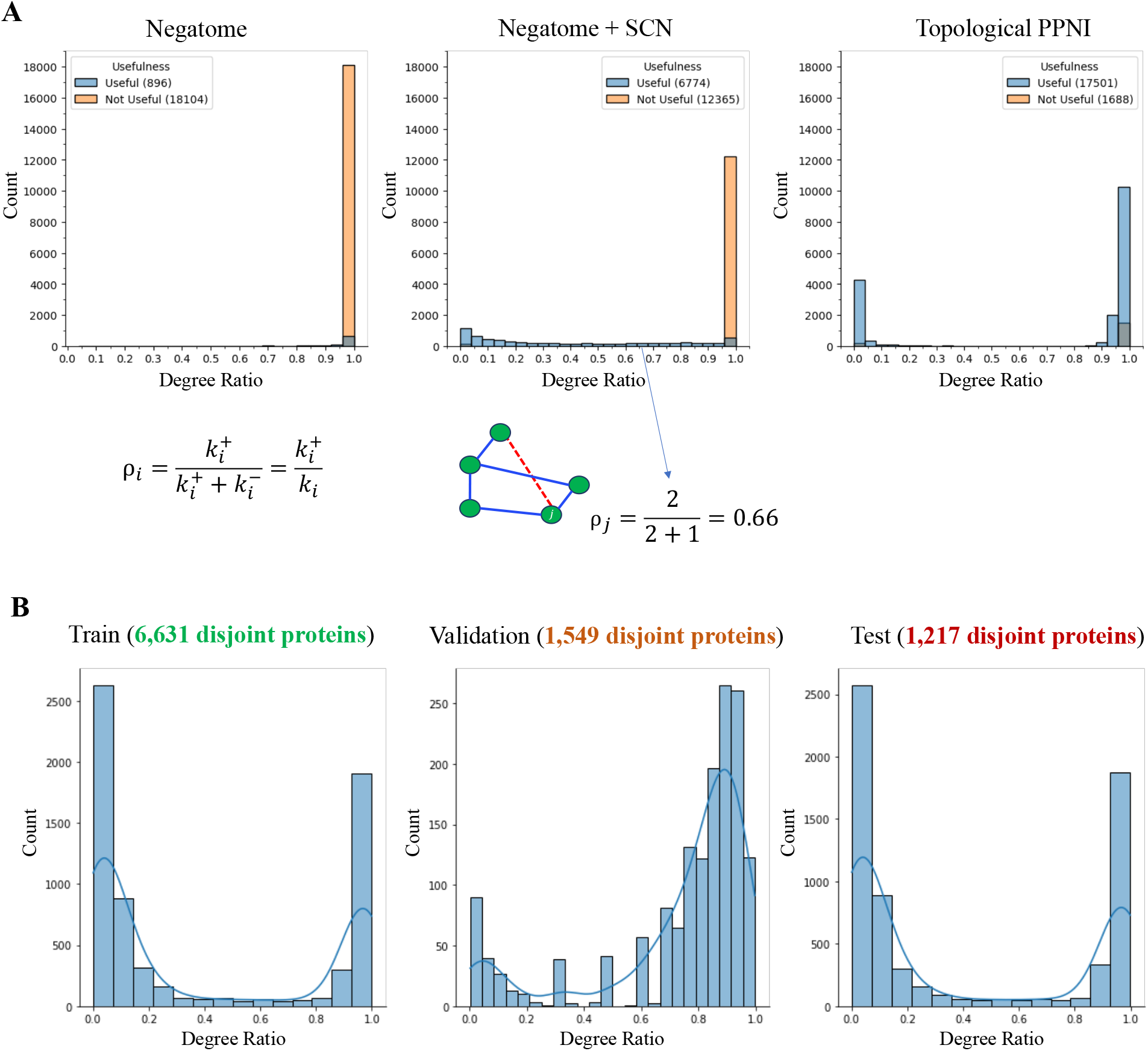
Removing annotation imbalance and topological shortcuts. **(A)** Combining the PPI databases and Negatome with only 2k PPNI samples creates a large annotation imbalance in the training data. The majority of proteins only have positive samples and lack negative examples. This imbalance is quantified by degree ratio 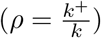 [5], which is measured by the ratio of positive annotations (*k*^+^) to total annotations (*k* = *k*^+^ + *k*^*−*^) associated with a protein. Subcellular Compartmental Negatives (SCN) introduces 778k PPNI examples, rectifying annotation imbalance partially. Finally, topological PPNI fixes the issue of annotation imbalance by introducing a substantial number (3 million) of non-interactions. We then filter the interactions and non-interactions involving the proteins having both positive and negative samples for training and testing of UPNA-PPI. Removing proteins with only positive or negative samples helps UPNA-PPI circumvent learning shortcuts related to predicting that some proteins always interact or do not interact. **(B)** Using only topological PPNI we remove annotation imbalance from all train, validation, and test data of UPNA-PPI. Here, we visualize the degree ratio distributions in train, validation, and test datasets for the first fold.

**Figure S5:**
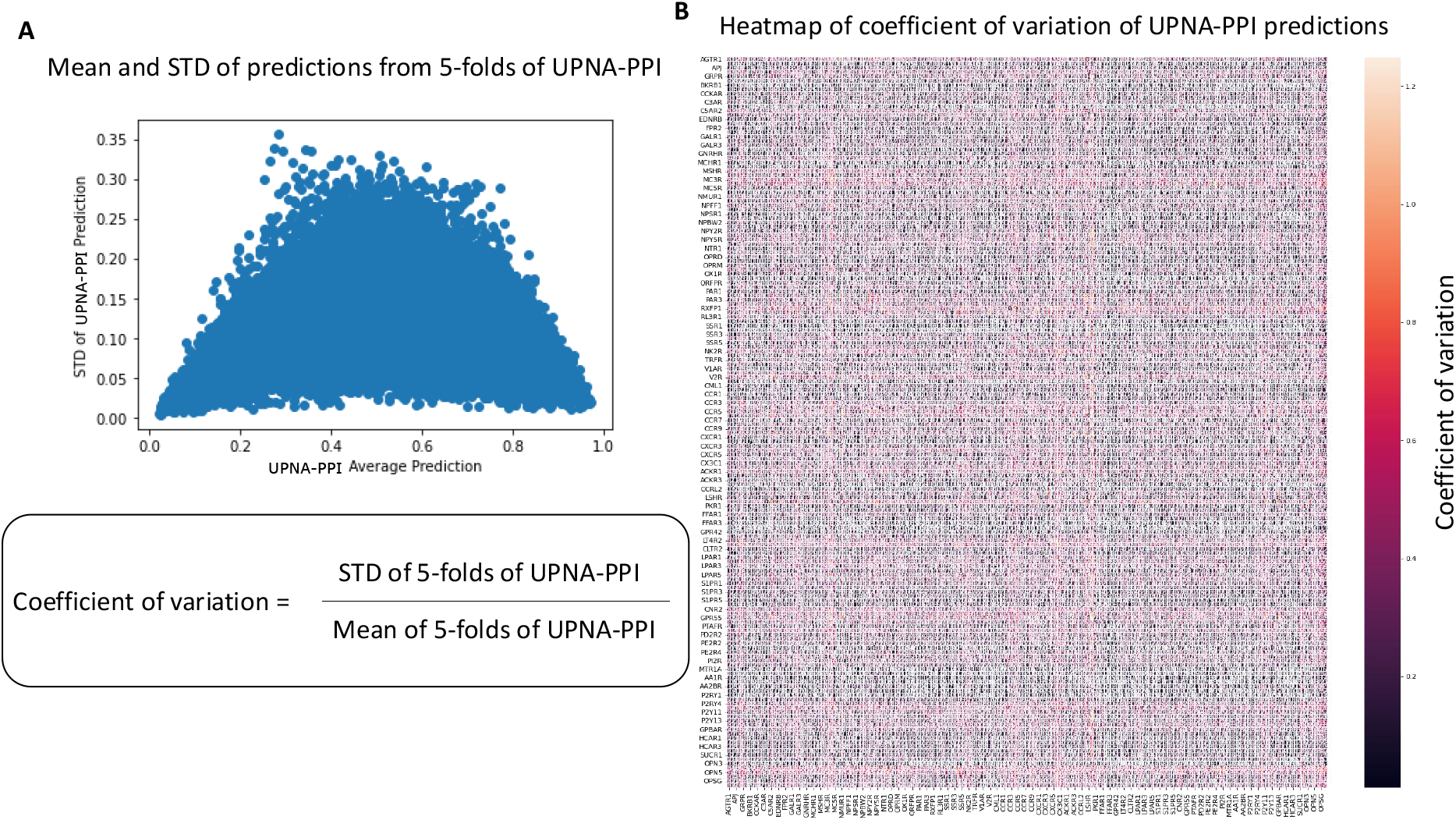
Coefficient of variation. **(A)** We plot the standard deviation (STD) vs. mean of the predictions from the 5-folds of UPNA-PPI for GPCR proteins. We observe that the STD is lower for the predictions near 0 and 1, indicating that UPNA-PPI models from 5-folds agree upon the top and the bottom predictions. Whereas, the predictions in between have higher STD. Thus, we define coefficient of variation of UPNA-PPI predictions as the ratio of the STD and mean from the 5-folds. Lower value of this coefficient suggests lower STD with high average, this high confidence interaction predictions. **(B)** We plot the heatmap of the coefficient of variation from UPNA-PPI predictions for 170 GPCR proteins. The darker dots correspond to the high confidence interaction predictions.

## Footnotes

* Precision at top K predictions with K=100

